# A comprehensive temporal proteome and phosphoproteome dataset of Y1 adrenocortical cells under FGF2 stimulation and SOS inhibition

**DOI:** 10.64898/2026.10.19.759539

**Authors:** Fabio Montoni, Toshiki Hanada, Doi Kentaro, Rosangela Aparecida Moreira Wailemann, Jun Adachi, Marcelo da Silva Reis, Yayoi Natsume-Kitatani

**Affiliations:** Center of Toxins, Immune-Response and Cell Signaling (CeTICS), Butantan Institute, São Paulo, 05503-900, Brazil; Cell Cycle Laboratory, Butantan Institute, São Paulo, 05503-900, Brazil; Bioinformatics Graduate Program, University of São Paulo, São Paulo, 05508-900, Brazil; Laboratory of Artificial Intelligence and Inference in Complex Data (Recod.ai), Institute of Computing, University of Campinas, Campinas, 13083-852, Brazil; AI Center for Health and Biomedical Research (ArCHER), National Institutes of Biomedical Innovation, Health and Nutrition, Osaka, 566-0002, Japan; Laboratory of Proteomics for Drug Discovery, Center for Drug Design Research (CDDR), National Institutes of Biomedical Innovation, Health and Nutrition, Ibaraki, 567-0085, Japan

**Keywords:** Phosphoproteomics, Y1 Adrenocortical Cell, Cancer Research, Cell Signaling, Mass Spectrometry

## Abstract

The Y1 mouse adrenocortical tumor cell line is a classic model for cancer studies and ACTH-mediated steroidogenesis. The KRAS gene amplification and SOS-mediated activation sustain elevated ERK levels, which drive the tumor phenotype of this cell line. In addition, the mitogen FGF2 induces atypical cytotoxic stress and consequent cell-death rather than proliferation in this system, imposing a strong perturbation on KRAS-ERK signaling dynamics. Recent research from our group revealed that SOS is not solely responsible for KRAS activation in the Y1 cells, indicating a more complex regulatory landscape of GEFs and GAPs. This motivated the generation of comprehensive time-resolved proteome (DIA) and TMT-phosphoproteome (DDA) datasets over a 0–30 min time course, including FGF2 stimulation alone or combined with the SOS1 inhibitor BI-3406, in the presence or absence of FBS, with three independent culture replicates per experimental condition using Q-Exactive™ and Thermo Lumos™ mass spectrometers, respectively. This dataset can enable latent mechanisms between the life and death switch caused by FGF2 signaling and the alternate stress response caused by impaired SOS-dependent activation. This analysis yielded 14,512 phosphosites and 8,786 protein groups, where 10,416 phosphosites and 111 protein groups showed differential abundance with statistical significance (limma, FDR < 0.05) in at least one of the experimental conditions. This dataset provides a versatile resource for downstream applications, which includes dynamic modeling such as ordinary differential equation–based frameworks, for which temporal datasets remain scarce.

## Background and Summary

The Y1 mouse adrenocortical tumor cell line is a classic model for cancer studies and ACTH (adrenocorticotropic hormone)-mediated steroidogenesis. The tumor phenotype is sustained by the combination of two factors: (1) KRAS (Kirsten rat sarcoma viral oncogene homolog) oncogene amplification (**George et al., 1985**), (2) with high KRAS-GTP basal levels (**Salotti et al., 2013**), mediated by the Guanine Exchange Factor (GEF) SOS (Son of Sevenless) (**Bos, Rehmann and Wittinghofer, 2007**). In addition to that, the Fibroblast Growth Factor 2 (FGF2), known for its proliferative role (**Armelin, 1973**), induces Y1 to atypical cytotoxic stress and consequent cell-death rather than proliferation. FGF2 impairs the cell cycle in G2/M, restraining the proliferation of the Y1 cell line via the ERK (extracellular signal-regulated kinase) pathway, leading to cell death (**Salotti et al., 2013**). In contrast to what is already established, our research group revealed that SOS is not solely responsible for KRAS activation in the Y1 cells, and the GEF RASGRP4 (RAS guanyl-releasing protein 4) was found essential for sustaining the previously described tumor phenotype in Y1 cells (**Montoni et al., 2026a**). This indicated a complex regulatory landscape of the GEFs and GTPase-Activating Proteins (GAPs), which warrants exploration.

This motivated the generation of a comprehensive time-resolved proteome (DIA) and TMT-phosphoproteome (DDA) that could enable the latent mechanisms between the life and death switch caused by FGF2 signaling and the alternate stress response caused by impaired SOS-dependent activation, with the use of the inhibitor BI-3406. The inhibitor binds to a pocket adjacent to His905SOS1, which disrupts the Tyr884SOS1–Arg73RAS interaction that stabilizes the SOS1–KRAS complex (**Hofmann et al., 2021**). The proteome (DIA) and TMT-phosphoproteome (DDA) were performed in a time course that ranged from 0 to 30 minutes under FGF2 stimulation or combined with SOS inhibitor, with the presence or absence of FBS stimulation, in three independent culture replicates using Q-Exactive™ and Thermo Lumos™ mass spectrometers, respectively. This analysis yielded 14,512 phosphosites and 8,786 protein groups, where 10,416 phosphosites and 111 protein groups showed differential abundance with statistical significance (limma, FDR < 0.05) in at least one of the tested conditions. This dataset provides a versatile resource for downstream applications, which includes dynamic modeling such as ordinary differential equation– based frameworks, of which time course-based datasets remain scarce (**Almquist et al., 2014; Wittig et al., 2014; Du et al., 2016**). We believe this dataset is a relevant resource for studying cancer, specifically focusing on the MAPK (mitogen-activated protein kinase) signaling pathway.

## Methods

### Experimental Design

For each replicate, 1×106 Y1 cells were plated in the 15 cm2 dishes (IWAKI, Japan, Cat# 3143-225). The next day, the synchronization in the phase G0/G1 by serum starvation was performed by adding FBS-free DMEM (fetal bovine serum-free Dulbecco’s Modified Eagle Medium) for 48h. The experimental design comprised three treatment conditions: (1) control, without stimulation; (2) stimulation with 10 ng/mL fibroblast growth factor 2 (FGF2), prepared in-house according to Costa et al. (**Costa et al., 2008**); and (3) combined treatment with 10 ng/mL FGF2 and 0.5 nM of the SOS1 inhibitor BI-3406 (SOSi) (MedKoo Biosciences, United States, Cat# 207113). Each treatment condition was evaluated in the presence (+FBS) or absence (™FBS) of fetal bovine serum. Control samples were collected at 0, 5, 10, and 30 min, whereas FGF2- and SOSi + FGF2-treated samples were collected at 5 s, 5, 10, and 30 min. Each experimental combination included three independent culture replicates, yielding a total of 72 samples. The complete sample assignment can be seen in **Table 1 (Sample Assignment)**. A schematization can be observed in **Figure 1 (Experimental Design Schematization**.

**Table 1:** Sample identification and replicate assignment. Sample identification and replicate assignment: This table provides the correspondence between sample identification numbers and their respective samples and replicates, serving as a reference for identifying individual samples in the raw proteomics and phosphoproteomics datasets.

| **Condition** | **Time** | **Replicate 1** | **Replicate 2** | **Replicate 3** |
|---|---|---|---|---|
| **Control 0 min - FBS** | 0 min | 1 | 25 | 49 |
| **Control 0 min + FBS** | 0 min | 2 | 26 | 50 |
| **Control 5 min - FBS** | 5 min | 3 | 27 | 51 |
| **Control 5 min + FBS** | 5 min | 4 | 28 | 52 |
| **Control 10 min - FBS** | 10 min | 5 | 29 | 53 |
| **Control 10 min + FBS** | 10 min | 6 | 30 | 54 |
| **Control 30 min - FBS** | 30 min | 7 | 31 | 55 |
| **Control 30 min + FBS** | 30 min | 8 | 32 | 56 |
| **FGF2 5 sec - FBS** | 5 sec | 9 | 33 | 57 |
| **FGF2 5 sec + FBS** | 5 sec | 10 | 34 | 58 |
| **FGF2 5 min - FBS** | 5 min | 11 | 35 | 59 |
| **FGF2 5 min + FBS** | 5 min | 12 | 36 | 60 |
| **FGF2 10 min - FBS** | 10 min | 13 | 37 | 61 |
| **FGF2 10 min + FBS** | 10 min | 14 | 38 | 62 |
| **FGF2 30 min - FBS** | 30 min | 15 | 39 | 63 |
| **FGF2 30 min + FBS** | 30 min | 16 | 40 | 64 |
| **SOSi + FGF2 5 sec - FBS** | 5 sec | 17 | 41 | 65 |
| **SOSi + FGF2 5 sec + FBS** | 5 sec | 18 | 42 | 66 |
| **SOSi + FGF2 5 min - FBS** | 5 min | 19 | 43 | 67 |
| **SOSi + FGF2 5 min + FBS** | 5 min | 20 | 44 | 68 |
| **SOSi + FGF2 10 min - FBS** | 10 min | 21 | 45 | 69 |
| **SOSi + FGF2 10 min + FBS** | 10 min | 22 | 46 | 70 |
| **SOSi + FGF2 30 min - FBS** | 30 min | 23 | 47 | 71 |
| **SOSi + FGF2 30 min + FBS** | 30 min | 24 | 48 | 72 |

**Figure 1.**
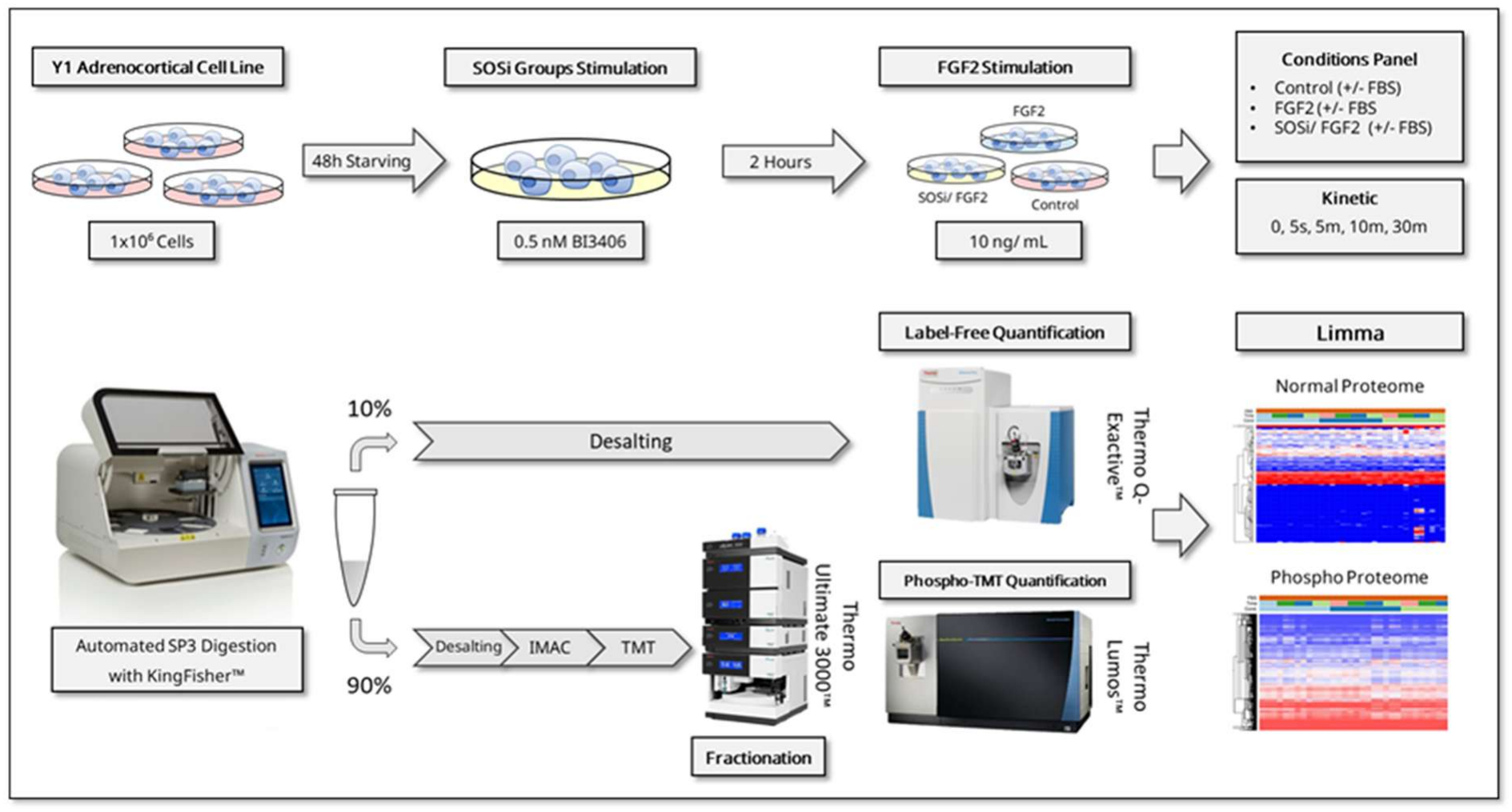
Experimental Design Schematization. Y1 mouse adrenocortical tumor cells were synchronized in G0/G1 by serum starvation and subjected to three experimental conditions: **(1)** unstimulated control; **(2)** stimulation with 10 ng/mL fibroblast growth factor 2 (FGF2); or **(3)** combined treatment with 10 ng/mL FGF2 and 0.5 nM of the SOS1 inhibitor BI-3406 (SOSi). Each condition was evaluated in the presence or absence of fetal bovine serum (FBS). Control samples were collected at 0, 5, 10, and 30 min, whereas FGF2- and SOSi + FGF2-treated samples were collected at 5 s, 5, 10, and 30 min. Each experimental combination included three independent culture replicates. Cell lysates were processed for proteomic and phosphoproteomic analyses, with the majority of the material allocated to phosphopeptide enrichment and the remainder to total proteome analysis. The phosphoproteome and proteome datasets were acquired using Orbitrap-based mass spectrometry platforms. Statistically significant changes were identified using linear models implemented in the limma framework.

### Cell Culture

The Y1 murine adrenocortical carcinoma cell line was obtained from ATCC. The cell was cultured in the Cell-IQ™ culture chamber (PHCBI, Japan, Series 5.8 cu.ft.) at 37 °C with 5% CO2 atmosphere in 225 cm2 Cell Culture Flasks (IWAKI, Japan, Cat# 3143-225) with Dulbecco’s modified Eagle’s media (Nacalai Tesque, Japan, Cat# 08458-45), supplemented with 10% Fetal Bovine Serum (Invitrogen, United States, Cat# 10437028) and Penicillin/ Streptomycin 1% (Gibco, United States, Cat# 15140-163). When necessary, the cell passage or separation for cell counting was performed by removing the culture media and washing with PBS-A (phosphate-buffered saline A) (Invitrogen, United States, Cat# 10437028), and after its removal, the addition of 1:300 trypsin (Thermo, United States, Cat# 25300054) for 3 minutes. For the cell counting, the Countess™ (Invitrogen, United States, Countess™ I) was used by analysing a 1:1 mixture of cell suspension and 0.4% Trypan Blue (Gibco, United States, Cat# 15250-061), taking into account only the cell amount assigned as viable to determine the cell volume for the experimentations.

### Protein Extraction

Immediately after the aforementioned stimulations, the dishes were placed in ice and washed twice with 5 mL of ice-cold PBS A (no inhibitors) and subsequently harvested with 1 mL of ice-cold PBS A plus inhibitors (10 mM EDTA, PhosSTOP™ (Roche, Switzerland, Cat# 4906845001) and complete™ (Roche, Switzerland, Cat# 11873580001), pH 7.4), by using a plastic cell scraper. The cell suspension was transferred into a pre-cooled 1.5 mL Proteosave™ tube (Sumitomo Bakelite, Cat# MS-4215MZ) and centrifuged at 300 x g for 5 minutes at 4°C. The supernatant was carefully aspirated, and the samples were flash frozen in liquid nitrogen and stored in the -80 freezer until the next applications. In sequence, the cell pellet was resuspended in 300 μL of the PTS buffer (12 mM SDC, 12 mM SLS, 100 mM Tris-HCl, 10 mM EDTA, PhosSTOP™ and cOmplete™, pH 6.8), without pipetting up and down, and vortexed vigorously until there was no cell debris. The samples were incubated on a heat block at 95ºC for 5 minutes and subsequently placed on ice for 2 minutes. After that, the samples were sonicated in the BIORUPTOR® II (Cosmo Bio, Japan, BR2024A) in a cycle of 30 minutes, shifting between “On” and “Off” every 30 seconds on “High” mode. After sonication, the samples were checked to ensure there was no viscosity. In sequence, the tubes were centrifuged at 20,000 x g at 4ºC for 10 minutes. The supernatant was transferred to a new Proteosave™ tube. For the protein quantification, 5 μL of each sample was aliquoted separately to a 0.5 mL Safe-Lock Tube (including the lysis buffer for blank) and diluted 10 times by Milli-Q® water for protein quantification, performed with the DC Protein Assay™ Kit II (BioRad, United States, Cat# 5000112), by following strictly the manufacturer’s recommendations and subsequently read in the spectrophotometer iMark™ (Biorad, United States, iMark™ Microplate Reader) at 360 nm absorbance and the protein estimation curve performed by the Microplate Manager® (V.6.1) software. After that, the samples were flash frozen in liquid nitrogen and stored in the -80 freezer until the next application.

### Automated SP3 Digestion

This procedure was performed according to Muraoka et al. (**Muraoka et al., 2022**), in an automated protocol using the Kingfisher (Thermo, United States, APEX) system. To prepare the beads, prior to the experimentation, 1:1 (v:v) of 50 μg/μL of SP3 beads solution, composed of SP3 beads A (GE Healthcare, United States, Cat# 45152105050250) and SP3 beads B (GE Healthcare, United States, Cat# 65152105050250), was washed with 200 *µ*L Milli-Q® water, placed in the magnetic rack (Tamagawa, Japan, Cat# TAB4899N12) for 2 minutes and removed the water twice. After that, the 200 *µ*L Milli-Q® water was added. In sequence, for each sample, 100 *µ*g in 100 *µ*L of protein was carefully loaded in a Deepwell 96-well Plate (Thermo, United States, Cat# 97002570), and sequently, for reduction and alkylation a mixture of 4 *µ*L 0.25 M TCEP (Fujifilm WAKO, Japan, Cat# 208-21541) and 4 *µ*L 0.5 M IAA (iodoacetamide) (Nacalai Tesque, Japan, Cat# 19302-54) was added respectively at 37ºC for 60 minutes in the dark. After that, in the Kingfisher, the beads were mixed with samples for 30 minutes at medium intensity for binding the proteins to the beads, then sequentially washed with ACN (acetonitrile) (Kanto Pure Chemical, Japan, Cat# 01033-76), 70% EtOH (ethanol) (Fujifilm WAKO, Japan, Cat# 057-00451) for 3 minutes twice, followed by the beads collection. In sequence, the beads were transferred to the plate containing 100 *µ*L Digestion Buffer (50 mM HEPES (CST, United States, Cat# 44686S), Trypsin (1:50) Fujifilm WAKO, Japan, Cat# 202-20081), Lys-C (1:50) (Fujifilm WAKO, Japan, Cat# 121-02541). After that, the plate was incubated overnight (16 hours) at 37ºC. The next day, the samples were transferred to a plate that contained 100 *µ*L Milli-Q® water and mixed for 4 minutes to disassociate the beads from the digested peptides. Lastly, the samples were acidified (pH <2) with 20 *µ*L TFA (Fujifilm WAKO, Japan, Cat# 204-02743) to the final concentration of 1%. Immediately after the acidification, the samples were desalted by using OASIS HLB 30 mg cartridges (Waters, United States, Cat# WAT094225) loaded in a Pressure +48 Biotage® Manifold (Biotage, Sweden, Pressure +48). The cartridges were conditioned with 1.5 mL ACN and were loaded at “Strong” flow twice. After that, 1 mL ACN 60%, TFA 0.1%, followed by 1 mL TFA 0.1% in “Strong” mode to condition the column. Next, all the peptides were added into the column at “Weak” flow through mode, allowing slow binding to the column. To wash the column, 1 mL TFA 0.1% was loaded at “Weak” flow through mode twice. After that, the flow through mode was changed to “Strong” to remove residual sorbent or contaminants. Then the new 2.5 mL Proteosave tubes were assembled to the Pressure +48 Biotage® Manifold, and the samples were eluted with 0.4 mL ACN 60%, TFA 0.1% in “Weak” flow through mode, followed by a change to “Strong” mode, to allow the collection of all peptides without loss. From this step on, 10% of the eluate was separated for the DIA proteome, whereas 90% was separated for the DDA phosphoproteome. Lastly, the samples were dried in the Speedvac for 45 minutes in LC/MS/MS vials, with no heating function, and stored in the -80ºC freezer until the next application. The proteome samples were resuspended with 40 *µ*L of the LC/MS/MS Solution (2% ACN, 1% TFA) and the samples were stored at 4ºC degrees for 16 hours before the DIA proteome mass spectrometry.

### Phosphoproteome Enrichment with Fe-IMAC

This procedure was performed according to Adachi et al. (**Adachi et al., 2022**) and Gunji et al. (**Gunji et al., 2023**). In order to prepare the IMAC resin, the 25 mL Filtration Columns (Biotage, Sweden, Cat # 120-1026-E) were attached to a Pressure +48 Biotage® Manifold. The columns were washed with 10 mL of Milli-Q® water, and subsequently 4 mL of Probound™ Resin (Invitrogen, United States, Cat# R80115) was thoroughly vortexed, loaded into the columns, vacuum applied, and centrifuged at 3,000 × g for 1 minute to let them dry completely. Next, for the nickel elution, 12 mL of 50 mM EDTA, pH 8 (DOJINDO, Japan, Cat# 345-01865) was added and thoroughly mixed with the pipette, and then incubated for 3 minutes, and a vacuum was applied. After that, the EDTA solution was added again, thoroughly mixed with the pipette, centrifuged, and a vacuum was applied and centrifuged at 3,000 × g for 1 minute four times. After the nickel elution, the column was washed with 18 mL of Milli-Q® water, then 18 mL of 0.1% Acetic Acid (Fujifilm WAKO, Japan, Cat# 017-00251) was added, followed by applying vacuum and centrifugation at 3,000 × g for 1 minute. In sequence, the Fe binding procedure was performed by adding 18 mL of 100 mM Ferric Chloride (Nacalai Tesque, Japan, Cat# 19502-05) in 0.1% Acetic Acid in the Filtration Columns. This solution was mixed vigorously by pipetting 10 times, then once again by vacuum application and centrifugation at 3,000 × g for 1 minute. Finally, the excess of the Fe solution was washed with 18 mL of 0.1% Acetic Acid and 18 mL of 60% ACN, 0.1% TFA, followed again by vacuum application and centrifugation for both procedures. For the final washing step, the valve was closed, 10 mL of 60% ACN, 0.1% TFA was added, and the whole content of the resin was transferred three times. The final volume was adjusted to 36 mL with 60% ACN, 0.1% TFA. The resin was maintained at 4°C until the day of use. For the phosphopeptide enrichment with stage-IMAC, 1x Empore C18 disk (3M, United States, Cat # 2215) was pressed into a pipette tip, to create a standard stage tip, as described in Adachi et al. (**Adachi et al., 2016**). After that, the IMAC Resin was vortexed vigorously for 10 seconds, and 200 *µ*L was added into the tip twice, and subsequently centrifuged at 2,000 × g for 3 minutes. The dried tip was immediately subjected to the phosphopeptide enrichment. Subsequently, 200 *µ*L of the peptide sample in 60% ACN, 0.1% TFA buffer was added to the Stage-IMAC tip, followed by centrifugation at 1,200 × g for 5 minutes twice. The columns were then washed with 200 *µ*L 60% ACN, 0.1% TFA buffer, then centrifuged at 2,000 × g for 3 minutes three times. In sequence, 200 *µ*L 0.1% TFA buffer was added and centrifuged at 2,000 × g for 3 minutes. For removing the non-phosphorylated peptides, 200 *µ*L of 0.1% TFA and 1% phosphoric acid was added and centrifuged at 1,200 × g for 5 minutes. This procedure was followed by a final wash with 200 *µ*L of 0.1% TFA and centrifugation at 2,000 × g for 3 minutes. For elution, the Stage-IMAC tip was transferred to a new ProteoSave tube. Then, 35 μL of 60% ACN, 0.1% TFA was added to elute phosphorylated peptides, followed by centrifugation at 1,500 × g for 3 minutes. The eluates were concentrated using a SpeedVac for 45 minutes, yielding enriched phosphorylated peptides for the Phospho-TMT Labeling procedure.

### Phospho-TMT Labeling and High-pH fractionation

Each sample was dissolved in 10 *µ*L of 100 mM TEAB (Sigma, United States, Cat# T7408), mixed at 1,200 rpm for 5 minutes in the EYELA tube mixer (EYELA, Japan, CM-1000), then centrifuged at 10,000 x g for 1 minute at room temperature. After this procedure, the TMTpro™ reagent (Thermo Scientific, United States, #A44520 [Lot# YB355487]) was vortexed, spun down, and immediately placed on ice until the moment of use. After that, 4 *µ*L of the reagent was added according to **Table 2 (Batch Assignment)**, and tubes were gently vortexed, spun down, and incubated at room temperature for 1 hour. To quench the reaction, 1 *µ*L of 5% hydroxylamine (Fujifilm WAKO, Japan, Cat# 088-07221) in 100 mM TEAB was added to each sample, then once more vortexed, spun down, and incubated for 15 minutes at room temperature to allow the reaction. Finally, the tubes were centrifuged at 10,000 × g for 1 minute, and all labeled samples were combined into a single tube, including the reference sample in each batch (1:1 of every sample), always using the same pipette tip, and then the tubes were dried at speedvac. For the cleanup procedure, 2x C18 + 1x Empore SCX (3M, United States, Cat# 2251) disks were mounted as described in Adachi et al (**Adachi et al., 2016**). The previously dried samples were dissolved in 50 *µ*L of Buffer U (8M Urea (Fujifilm WAKO, Japan, Cat# 219-00175), 20% TFA) and mixed at 1,200 rpm for 5 minutes in EYELA. In sequence, the samples were centrifuged at 20,000 × g for 20 minutes at 4°C. During this period, the tips were conditioned with 30 *µ*L and spun at 2,000 x g for 1.5 min in the following sequence: methanol, Buffer SB (0.5% TFA, 80% ACN), Buffer SA (0.5% TFA, 5% ACN), Buffer S500 (2M NH4AcO (Fujifilm WAKO, Japan, Cat# 015-02832), 30% ACN), and Buffer SA. After that, each sample was loaded in 30 *µ*L aliquots and centrifuged at 2,000 × g for 1.5 minutes. For the sample washing procedure, 50 *µ*L was added and spun at 2,000 x g for 3 minutes in the sequence: Buffer U, Buffer SA, Buffer SB, Buffer SA, and Buffer SB. Finally, the samples were eluted with 35 *µ*L and centrifuged at 2,000 × g for 2 minutes and dried at speedvac. Each batch of the TMT-Labeled samples was separated into seven fractions in a high pH gradient using the Ultimate™ 3000 UHPLC (Thermo, United States, Ulti-Mate™ 3000 UHPLC system) according to Shoji et al. (**Shoji et al., 2024**) and Gunji et al. (**Gunji et al., 2024**). The solutions were: Solution A (ACN 5%, 0.5 mM ABC, pH 9.2) and Solution B ACN 60%, 0.5 mM ABC, pH 9.2). The gradient was: 5% B to 25% over the first 4 minutes, then rising steadily to 60% by 30 minutes. After the fractionation procedure, each sample was immediately dried in the SpeedVac, and subsequently resuspended with 10 *µ*L of the LC/MS/MS solution, vortexed carefully in the EYELA 1200 RPM for 5 minutes, and then submitted to the DDA phosphoproteome mass spectrometry.

**Table 2:** TMT-based experimental batch and sample assignment. Overview of sample allocation across TMT experimental batches, with batch assignments randomized to minimize potential batch-related biases. Sample identification numbers correspond to those defined in Table 1, enabling consistent sample tracking across experimental batches.

|  | **Sample Number** |  |  |  |  |  |  |  |  |  |  |  |  |  |  |  |
|---|---|---|---|---|---|---|---|---|---|---|---|---|---|---|---|---|
| **Batch 1** | 1 | 16 | 31 | 46 | 61 | 2 | 17 | 32 | 47 | 62 | 3 | 18 | 33 | 48 | 63 | Ref. |
| **Batch 2** | 4 | 19 | 34 | 49 | 64 | 5 | 20 | 35 | 50 | 65 | 6 | 21 | 36 | 51 | 66 | Ref. |
| **Batch 3** | 7 | 22 | 37 | 52 | 67 | 8 | 23 | 38 | 53 | 68 | 9 | 24 | 39 | 54 | 69 | Ref. |
| **Batch 4** | 10 | 25 | 40 | 55 | 70 | 11 | 26 | 41 | 56 | 71 | 12 | 27 | 42 | 57 | 72 | Ref. |
| **Batch 5** | 13 | 28 | 43 | 58 | 14 | 29 | 44 | 59 | 15 | 30 | 45 | 60 |  |  |  | Ref. |

### DIA Proteome Mass Spectrometry

This methodology was performed with the Q Exactive™ mass spectrometer coupled with an UltiMate 3000 NanoLC system and an HTC-PAL autosampler. For each sample, 200 ng was loaded onto an analytical column (75 µm × 30 cm, packed in-house with ReproSil-Pur 120 C18-AQ, 1.9 µm resin (Dr. Maisch, Germany, Cat. # r119.aq.)). The gradient was set at a flow rate of 280 nL/min using a 110-minute gradient from 5% to 30% solvent B in 105 minutes, then 95% solvent B for 5 minutes. For the DIA, the gasphase fractionation GPF-DIA acquisition of a pooled sample for library generation, full mass spectra were acquired with the following parameters: a resolution of 120,000, an automatic gain control (AGC) target of 300%, and a maximum injection time in auto mode. The five GPF-DIA runs collectively covered 499-861 m/z (499-573, 571-645, 643-717, 715-790, and 787-861 m/z). MS2 spectra were collected with the following parameters: a 2-m/z isolation window at 60,000 resolution, AGC target of 3,000%, a maximum injection time in auto mode, and an NCE of 30.

### DDA Phosphoproteome Mass Spectrometry

This methodology was performed in data-dependent acquisition (DDA) mode using an Orbitrap Fusion™ Lumos™ Tribrid™ mass spectrometer (Thermo Fisher Scientific) coupled to an UltiMate™ 3000 NanoLC system and an HTC-PAL autosampler. The entire phosphopeptide sample was loaded onto an analytical column (75 μm × 30 cm) packed in-house with ReproSil-Pur C18-AQ resin (1.9 μm; Dr. Maish, Germany). Peptides were separated at a flow rate of 280 nL/min using a 180-minute linear gradient from 5% to 30% solvent B over 175 minutes, followed by a wash at 95% solvent B for 5 minutes. Full MS scans were acquired in the Orbitrap at a resolution of 120,000 over an m/z range of 375–1500, with an AGC target of 4.0 × 10^5^ and a maximum injection time of 50 ms. The default charge state was set to 2, and data were acquired in positive ion mode. Data-dependent MS/MS acquisition was performed using a Top12 method, with precursor ions isolated in the quadrupole using a 0.7 m/z isolation window and fragmented by higher-energy collisional dissociation (HCD) at a normalized collision energy of 32%. MS2 spectra were acquired in the Orbitrap at a resolution of 50,000, with an AGC target of 1.0 × 10^5^ and a maximum injection time of 105 ms. Dynamic exclusion was enabled with a duration of 30 seconds.

### DIANN

For the DIA proteome samples, the DIA-NN (ver. 1.8.2) (**Demichev et al., 2020**) analysis was conducted using the GPF spectral library generated previously, and the FASTA databases for the main search and contaminants removal were the same used in the Maxquant DDA analysis. For the parameters, the precursor ions were generated by performing an in silico digestion using Trypsin/P with up to two missed cleavages. Carbamidomethylation of cysteine (+57.02146 Da) was set as a fixed modification, while oxidation of methionine (+15.99491 Da) was considered a variable modification, with a maximum of two variable modifications per peptide. The length of peptides was fixed between 7 and 30 amino acids, with precursor charge states ranging from 2 to 4 and an m/z range of 300 to 1800. The mass accuracy for precursor ions was set to 10 ppm, while MS1 accuracy was set to 4 ppm. Isotopologues were included in the analysis, and a match-between-runs (MBR) strategy was enabled. Likely interferences were removed, and a single-pass mode neural network classifier was applied for data processing. Protein inference was performed using the first gene name, and quantification was carried out using a robust LC normalization strategy without decoys. Cross-run normalization was RT-dependent, and the peptide library was generated from the provided FASTA database. For precursor filtering, fragment ions were selected within an m/z range of 200 to 1800, and peptides with missed cleavages were limited to two. A precursor false discovery rate (FDR) of 1.0% and a protein group FDR of 1.0% were applied.

### MaxQuant

For the analysis of the Phospho-TMT samples, the program MaxQuant (version 2.6.2.0) (**Tyanova, Temu e Cox, 2016**) was used. For all the cases, the same parameters were applied. For the database, the *Mus musculus* Swiss Prot from Uniprot (downloaded in June 2024) and the MaxQuant default contaminants.fasta database were used. All the fractions were joined in the same experiment. The global search parameters were: Typsin and Lys-C, up to two missed cleavage sites, carbamidomethylation of cysteine (+57.02146 Da) residues set as the fixed modification, and N-terminal protein acetylation (+42.01056 Da) and methionine oxidation (+15.99491 Da) set as variable modifications. The Reporter on MS2 TMT-16-Pro label was enabled, and the reporter ion isotopic distributions were used according to the lot # YB355487. For phosphoproteome parameters, the phosphorylation of serine, threonine, and tyrosine was added as a variable modification. The PIF (precursor ion fraction) filter was set at 50%. The search results were filtered to a maximum false discovery rate (FDR) of 0.01 at the protein, peptide-spectrum match (PSM), and posttranslational modification (PTM) site levels. Two or more unique/razor peptides were required for protein identification, and a ratio count of 2 or more for protein quantification. PTM sites with a measured localization probability >0.75 were considered confident phosphosites. No normalization was set.

### Preprocessing and Bayesian PCA-based imputation (BPCA)

Raw data tables were filtered to remove decoy sequences (Reverse) and potential contaminants. Proteins with more than 30% missing or zero values across samples were discarded to ensure robust quantification. The remaining values were log2-transformed, with zeros replaced by NaN to avoid artifacts in subsequent analysis. Missing values were then imputed using Bayesian PCA with automatic relevance determination (BPCA-ARD), implemented through the bpca-impute package (**Oba et al., 2003**) using a Python adaptation (**Kitamoto et al., 2023**). The imputation was performed using 35 latent components (‘n_PCs=35’ for phosphoproteome and ‘n_PCs=5’ for proteome), strict convergence thresholds (‘eps=1e-4’), and a maximum of 1000 iterations (‘itr_max=1000’) to ensure stable recovery of missing intensities without excessive smoothing. Data handling and transformation steps were performed using pandas (**McKinney and others, 2011)** and numpy (**Harris et al., 2020**).

### ComBat Batch Effect Correction

Batch correction was applied to the BPCA-imputed data using the pycombat implementation (**Behdenna et al., 2023**) of the ComBat algorithm Experimental condition was included as a covariate in the model to account for biological variation during batch correction. The sample assignments for each TMT batch are provided in **Tables 1 (Sample Assignment) and 2 (Batch Assignment)**.

### Feature mean-centering

After batch correction, feature intensities were mean-centered across samples to set the average value of each feature to zero, without scaling the variance. This procedure was performed using StandardScaler from scikitlearn (**Pedregosa et al., 2011**) with centering enabled and standardization disabled. The resulting centered matrix was ready for downstream analyses.

### Dimensionality Reduction Methods

Principal Component Analysis (PCA) was performed using the scikit-learn implementation (**Pedregosa et al., 2011**), where the first two components were used to generate scatterplots highlighting group separation. Uniform Manifold Approximation and Projection (UMAP) was computed with the umap-learn package (**McInnes, Healy e Melville, 2018**).

### Differential protein/ phosphorylation with Limma

Differentially abundance proteins/ phosphorylated sites were identified using the limma (**Ritchie et al., 2015**) package in R. Log-transformed intensity values were used to build a design matrix representing the experimental conditions and time points. A single linear model was fitted across all samples, and empirical Bayes moderation was applied to obtain moderated t-statistics, p-values, and log2 fold changes (logFC) for each phosphosite. Differential protein/ phosphorylation abundance was evaluated through predefined contrasts comparing treated and control groups at each time point. An absolute log2 fold change greater than log2(1.5) and an adjusted p-value below 0.05 were considered significantly regulated, and were classified as up- or downregulated according to the direction of the logFC. The contrasts assignment can be observed in **Table 3 (Limma Contrasts Assignment)** and the results in **Supplementary Table 1 (Proteome Limma Analysis)** and **Supplementary Table 2 (Phosphoproteome Limma Analysis)**.

**Table 3.**
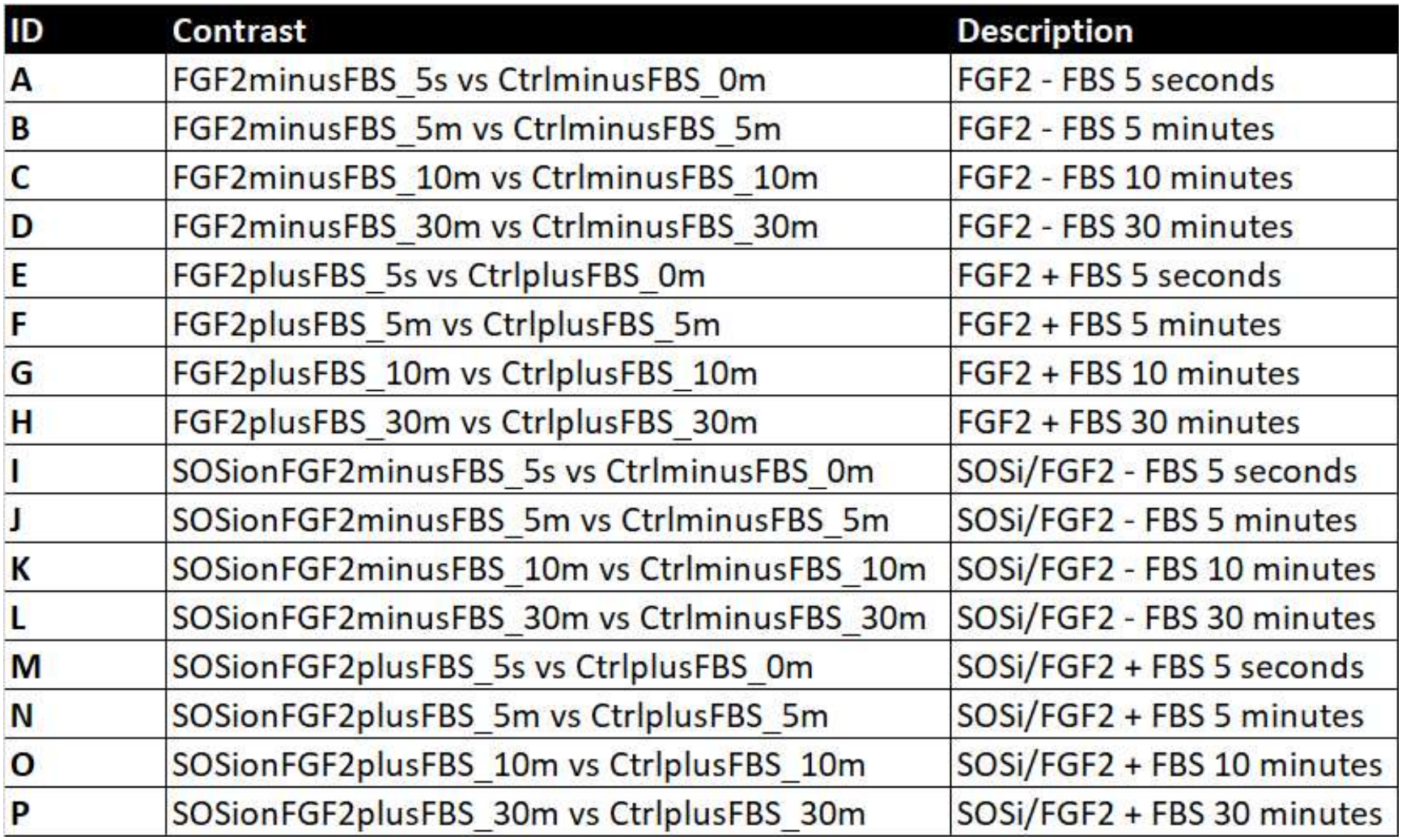
Limma contrast definitions. Summary of the pairwise comparisons between experimental conditions used to define the contrasts for differential expression analysis using Limma. Each contrast specifies the conditions compared to identify differences in protein and phosphoprotein abundance.

### Gene Enrichment

Over-representation analysis (ORA) was performed using WebGestalt (WEB-based GEne SeT AnaLysis Toolkit; https://www.webgestalt.org) (**Elizarraras et al., 2024**). Analyses were conducted for Mus musculus using the Gene Ontology Biological Process (GO-BP, non-redundant) annotation set. Protein identifiers were provided as UniProt Swiss-Prot IDs and mapped internally by WebGestalt to Entrez Gene identifiers. The reference set was defined as all protein-coding genes annotated in the mouse genome available in the WebGestalt platform. Categories containing fewer than 5 or more than 2,000 genes were excluded. Statistical significance was evaluated using the Benjamini–Hochberg false discovery rate (FDR) correction. Enrichment results were ranked according to adjusted p-values. The KEGG pathway enrichment was performed by using STRING (https://string-db.org/) enrichment tools (**Szklarczyk et al., 2024**).

### Data Visualization

Plots and graphical summaries were generated using matplotlib (**Hunter, 2007**) and seaborn (**Waskom, 2021**), with the scienceplots style (**Garrett, 2021**) applied. Heatmaps were generated using Morpheus web tool (https://software.broadinstitute.org/morpheus/). The kinome tree was created using Coral (**Metz et al., 2018**), by converting the mouse kinase names to *Homo sapiens*.

All the reagents, materials, buffers, plastics machinery, tools and software used in this research are enlisted in the **Supplementary Table 3 (Materials and Equipment List)**.

### Data availability

For this preprint version only the main text and supplementary figures are available. If you have interest in seeing the supplementary tables, raw data, processed tables and other information, please wait for the publication.

## Technical Validation

This experiment was conducted with the aim of comprehensively verifying the phosphorylation dynamics of the Y1 adrenocortical tumor cell. Under the effect of FGF2, whose cytotoxicity leads these cells to cell death by disturbing ERK levels, BI-3406, an SOS inhibitor, which may reveal alternative activation pathways of other GEFs, as well as the recently discovered RASGRP4, that showed as fundamental for the signalization of this cell. The experimental conditions comprised an unstimulated control, FGF2 stimulation (10 ng/mL), or combined treatment with FGF2 (10 ng/mL) and the SOS1 inhibitor BI-3406 (0.5 nM), each in the presence or absence of serum. Control samples were collected at 0, 5, 10, and 30 min, whereas FGF2- and SOSi + FGF2-treated samples were collected at 5 s, 5, 10, and 30 min. Each experimental combination included three independent culture replicates. Following stimulation, cells were rapidly collected on ice and lysed in the presence of protease and phosphatase inhibitors. Cell disruption was performed by sonication in the BIORUPTOR® II, and protein digestion was performed using an automated SP3 method with the KingFisher™ Apex system, which ensured homogeneous and reproducible lysis across all samples. Prior to the digestion, the samples pH were monitored (pH ∼7) to ensure the quality of the procedure. After completion, the samples were split in two: 10% was directed to the proteome and 90% was directed to the phosphoproteome enrichment. The Phosphoproteome enrichment was performed using an automated Fe^3+^-IMAC workflow, coupled to StageTip-based phosphopeptide purification, followed by TMT-labeling. Next, the phosphoproteome was fractionated with Offline high-pH reverse-phase peptide fractionation was conducted on an automated UltiMate™ 3000 UHPLC system equipped with a C18 column, producing concatenated peptide fractions for TMT-based LC–MS/MS analysis. Lastly, both proteome and phosphoproteome were desalted by using OASIS HLB 30 mg cartridges loaded in a Pressure +48 Biotage® Manifold, dried in SpeedVac and resuspended in LC/MS/MS solution for the mass spectrometry analysis.

### Spectral Data Acquisition

The proteome was analyzed using label-free data-independent acquisition (DIA) mass spectrometry on a Q Exactive™ mass spectrometer coupled to an UltiMate 3000 NanoLC system. The analysis yielded approximately 96,000 precursors and 8,786 protein groups. Peptide identification and protein quantification were performed using DIA-NN with an experimental GPF spectral library generated from a pooled sample comprising all experimental samples, using the Mus musculus reference FASTA file for protein sequence matching. Peptide and protein identifications were filtered at a false discovery rate (FDR) of ≤1%.

The phosphoproteome in turn was acquired in DDA (Data Dependent Acquisition) mode, using the Thermo Lumos™ mass spectrometer coupled with an UltiMate 3000 NanoLC system. Across all fractionated batches, a total of 2,637,504 MS/MS scans were submitted, of which 380,316 were identified. The MaxQuant summary reported 365,368 MS/MS identifications (SIL), 14,948 MS/MS identifications (PEAK), and 24,963 peptide sequences identified. A total of 3,627,307 isotope patterns were detected, of which 1,322,692 were sequenced and a total 14,512 phosphosites were identified. In the original experiment design, the reference sample containing 1:1 phosphopeptide mixture from all the samples (16th channel of each batch), was designated to be a normalizer of the multi batch conditions. However, our attempts to utilize the reference sample as a normalizer produced a cofounded pattern in all the samples. This limitation is commonly observed in multi-batch TMT-based experiments, showing that in some cases they can produce undesirable effects rather than assisting in the normalization step (**Herbrich et al., 2013; Ferretti et al., 2025**). Alternatively, a data-driven approach, which is widely applied to overcome normalizing issues (**Maes et al., 2016**), was applied to remove the batch effect and allow downstream analysis. This matter is addressed in detail in the section **Preprocessing and Normalization**.

### Preprocessing and Normalization

After acquiring the mass spectrometry data, both the proteome and the phosphoproteome underwent various filtering, normalization, and statistical techniques to ensure quality for downstream applications.

The proteome dataset originally contained 8,919 rows, corresponding to the number of protein groups identified. After the removal of contaminants, 8,786 rows remained. After log2 transformation, approximately 91.6% of the rows contained between 0 and 10% missing values (**Figure 2A**), and no retained row exceeded the predefined threshold of 30% zero values. In sequence, the boxplot analysis (**Figure 2C**) performed across the experimental conditions did not reveal significant variation among replicates or treatments. In addition, the boxplot whiskers exhibited minimal variation among replicates and treatments, indicating consistent data dispersion. Followed by that, the missing values were then imputed using Bayesian PCA with automatic relevance determination (BPCA-ARD) (**Oba et al., 2003**), implemented through the Python bpca-impute package (**Kitamoto et al., 2023**). The imputation was handled by using 5 latent components (‘n_PCs=5’), strict convergence thresholds (‘eps=1e-4’), and a maximum of 1000 iterations (‘itr_max=1000’).

**Figure 2.**
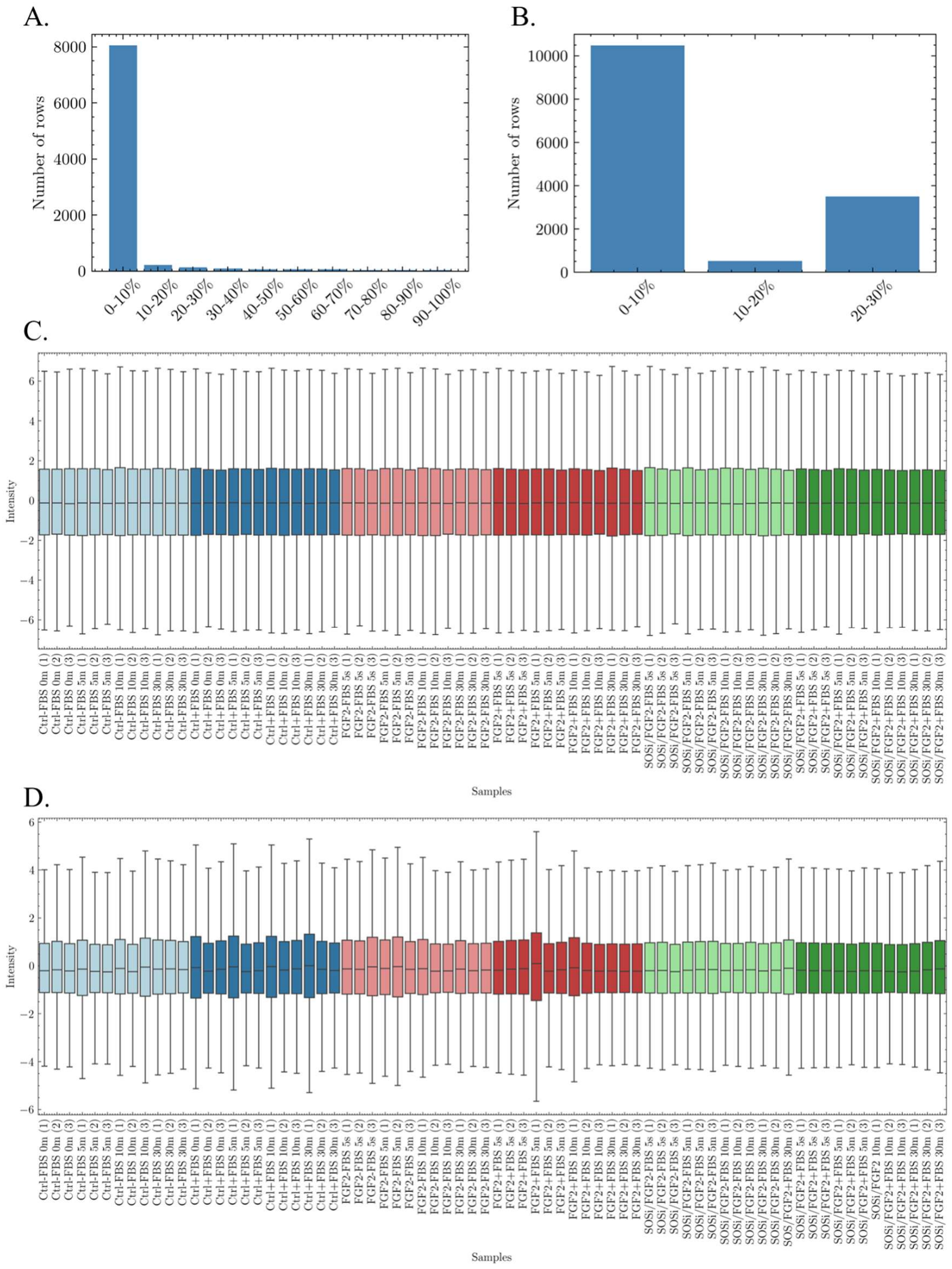
Preprocessing and Normalization. **A**: Distribution of zeros in the proteome matrix. Once lines presented only 10% max. zeros, no cutoff was performed. **B**: Distribution of zeros in the phosphoproteome matrix after lines >30% zeros were cutoff. **C**: Boxplot of distribution of proteome after applying the BPCA imputation method and mean centering. **D**: Boxplot of distribution of phosphoproteome after applying BPCA imputation method ComBat batch effect removal and mean centering. **Boxplot Legend**: **Light Blue**: Control - FBS, **Dark Blue**: Control + FBS, **Light Red**: FGF2 - FBS, **Dark Red**: FGF2 + FBS, **Light Green**: SOSi/FGF2 - FBS, **Dark Green**: SOSi/FGF2 + FBS.

The fractionated TMT phosphoproteome dataset contained 29,904 rows corresponding to the phosphosites in wide format, where each phosphosite has its value reported for one, two or three phosphorylations. The data were converted in Perseus from wide to long format using MaxQuant’s ‘Expand site table’ function, which tripled the number of rows from 29,904 to 89,712. After performing the removal of the reverse hits and contaminants, 88,689 rows remained. With this, a high proportion of missing values, with 68.0% of rows containing 90–100% missing values and only 11.8% containing 0–10%. However, this missingness largely reflects the structure of the data rather than true absence of measurements. As many sites lack values for the second or third phosphorylation state, this expansion introduced artificial missing values corresponding to non-existent phosphorylation, which were removed during filtering. After removing rows containing more than 30% zero values (**Figure 2B**), 14,512 rows were retained, representing approximately 50% of the initially identified phosphosites. The BPCA imputation was carried out by using 35 latent components (‘n_PCs=35’), strict convergence thresholds (‘eps=1e-4’), and a maximum of 1000 iterations (‘itr_max=1000’). Due to the aforementioned technical issues in the TMT-based experiments, the PyComBat (**Behdenna et al., 2023**) was applied in the data to remove batch effects. The boxplot analysis performed across the experimental conditions revealed slight differences in the size of Q1 and Q3 among replicates, with the data initially not centered. After applying ComBat correction and mean centering (**Figure 2D**), the data became comparable across replicates and treatments. Nonetheless, the boxplot whiskers demonstrated greater variation than previously observed in the normalized proteome, indicating increased data dispersion when compared to the proteome analysis.

To assess the impact of batch correction on the phosphoproteomic dataset, PCA and UMAP were used to visualize sample distributions before and after ComBat correction. The experimental conditions were distributed across the TMT batches, and experimental condition was included as a covariate in the pyComBat model to account for biological variation during batch correction. The sample assignments for each TMT batch are provided in **Tables 1 (Sample Assignment) and 2 (Batch Assignment)**. Before correction, PCA and UMAP revealed distinct clusters associated with batch identity (**Figures 3A and 3C**, respectively). After correction, these batch-associated patterns were no longer observed (**Figures 3B and 3D**). The first two principal components explained 60.51% of the total variance before correction and 55.59% after correction. When samples were colored according to experimental condition before correction (**Figures 3E and 3G**), the same batch-related groups remained apparent. After correction, some tendency for replicate samples to cluster together was observed in both PCA and UMAP projections (**Figures 3F and 3H**). However, the overall distribution remained complex, without a clear separation for all the experimental conditions.

**Figure 3:**
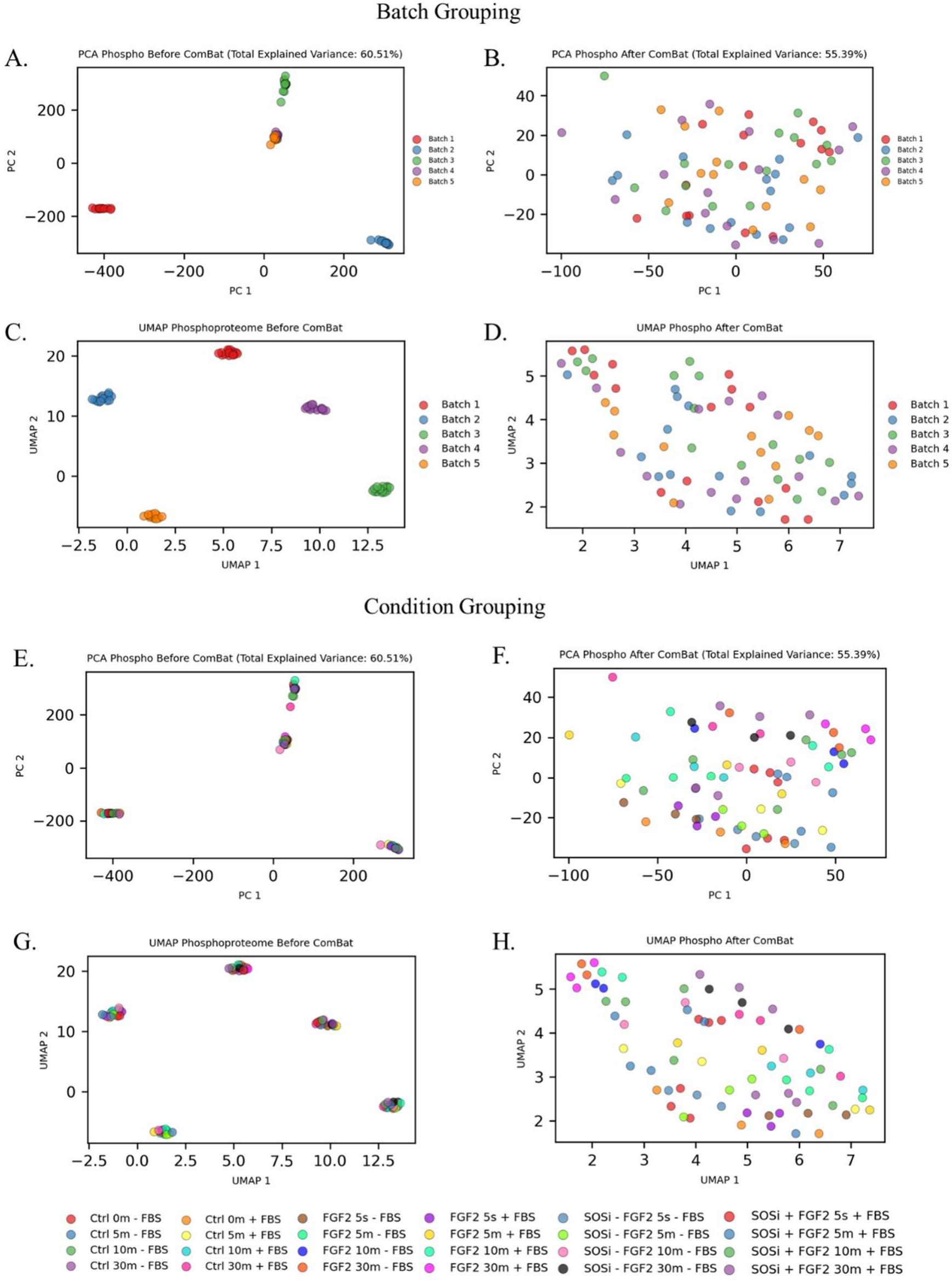
PCA and UMAP analyses of the phosphoproteomic dataset before and after ComBat batch effect correction. **A**: PCA before ComBat correction, colored by batch. **B**: PCA after ComBat correction, colored by batch. **C**: UMAP before ComBat correction, colored by batch. **D**: UMAP after ComBat correction, colored by batch. **E**: PCA before ComBat correction, colored by experimental condition. **F**: PCA after ComBat correction, colored by experimental condition. **G**: UMAP before ComBat correction, colored by experimental condition. **H**: UMAP after ComBat correction, colored by experimental condition. For PCA the first two principal components explained 60.51% and 55.59% of the total variance before and after ComBat correction, respectively.

Lastly, to confirm that all the preprocessing and normalization approaches have not altered the original data distribution, histogram analysis was performed for all the samples before and after the data imputation (**Figure 4**). **Figure 4A** shows the proteome normal distribution, and **Figure 4B** shows that all the imputed values remained within the overall distribution of the original data, suggesting that the imputation did not substantially alter the observed distributions. The same pattern can be observed for the phosphoproteome, where **Figure 4C** shows the normal distribution of the phosphoproteome data and **Figure 4D** shows that the number of the imputed values also fall within the original distribution of the data. In addition it can be also observed that the phosphoproteome data has a higher number of imputed values when compared to the proteome, common in TMT-based experiments.

**Figure 4:**
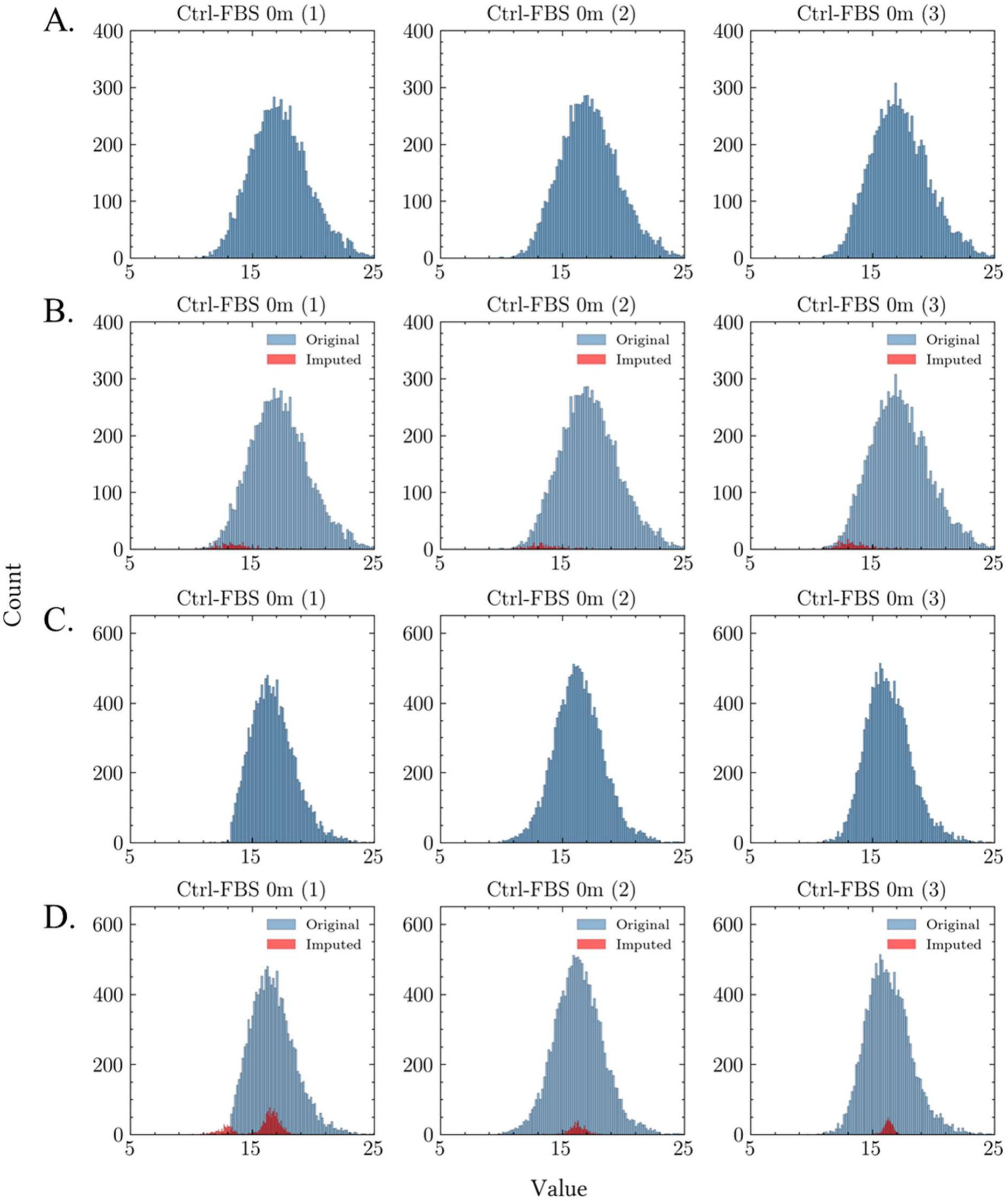
Example histograms of the frequency distribution of the samples across the processing procedures. This example features the control - FBS 0 minutes replicates to show how the preprocessing procedures does not affect/ change the original distribution of the data. **A**: Normal proteome unprocessed distribution. **B**: Normal proteome distribution after BPCA imputation methods. **C**: Phosphoproteome unprocessed distribution. **D**: Phosphoproteome distribution after BPCA imputation methods. The complete histograms can be accessed in the **Supplementary Material**.

### Proteins and Phosphosites Differential Abundance

After the preprocessing, the differential abundance of the proteins and phosphosites were assessed with the Limma package. For the proteome, most of the proteins remained stable in the 30-minute FGF2 stimulation timespan, having 111 proteins out of 8,786 with statistically significant changes in at least one of the conditions (**Figure 5A**). The major contrast can be observed in the SOSi/FGF2 treatments, where the inhibitor was added 2 hours prior to the FGF2 stimulation. For instance, proteins such as JUNB (JunB proto-oncogene) (UNIPROT ID: P09450), SERPINE1 (UNIPROT ID: P22777), CCN1 (cellular communication network factor 1) (UNIPROT ID: P18406) were found upregulated at SOSi/FGF2–FBS 5 min (2) and SOSi/FGF2 + FBS time points. Among the mentioned, JUNB is known for being an immediate early response transcription factor that is activated downstream of MAPK signaling and often associated with cell proliferation and differentiation responses (**Shaulian and Karin, 2002**), while SERPINE1, which plays an important role in cancer (**Shaikh et al., 2024**). Taken altogether, despite changes observed in the proteome analysis being only about 1.26% of the total, the statistically significant hits are strongly involved with cancer, in special the MAPK signaling, which controls the tumor phenotype observed in the Y1 cell. For the phosphoproteome, a complex pattern could be observed, having 10,416 differentially abundant phosphosites out of 14,512 in at least one of the conditions (**Figure 5B**), representing about 72% of the total phosphosites identified. The hierarchical clustering showed a complex pattern across all the experimental conditions. The analysis revealed a clear differentiation among the later time points under FGF2 stimulation. Specifically, samples from the FGF2–FBS 10 min (2–3) and FGF2–FBS 30 min (1–3) conditions clustered, and diverged from both the control and the early FGF2-stimulated samples. In addition, the FGF2+FBS 10 min (2–3) and FGF2+FBS 30 min (1–3) conditions displayed a comparable clustering pattern, indicating consistent proteomic changes at later time points. In contrast, the early FGF2 stimulations (5 s and 5 min.) remained closer to their respective controls, suggesting that major visible changes occur only after sustained exposure. The dynamic change of up and down-regulation of the phosphoproteome by condition can be observed in **Figure 6**. The result of the limma can be accessed on the **Supplementary Table 1 (Proteome Limma Analysis)** and **Supplementary Table 2 ( Phosphoproteome Limma Analysis)**.

**Figure 5:**
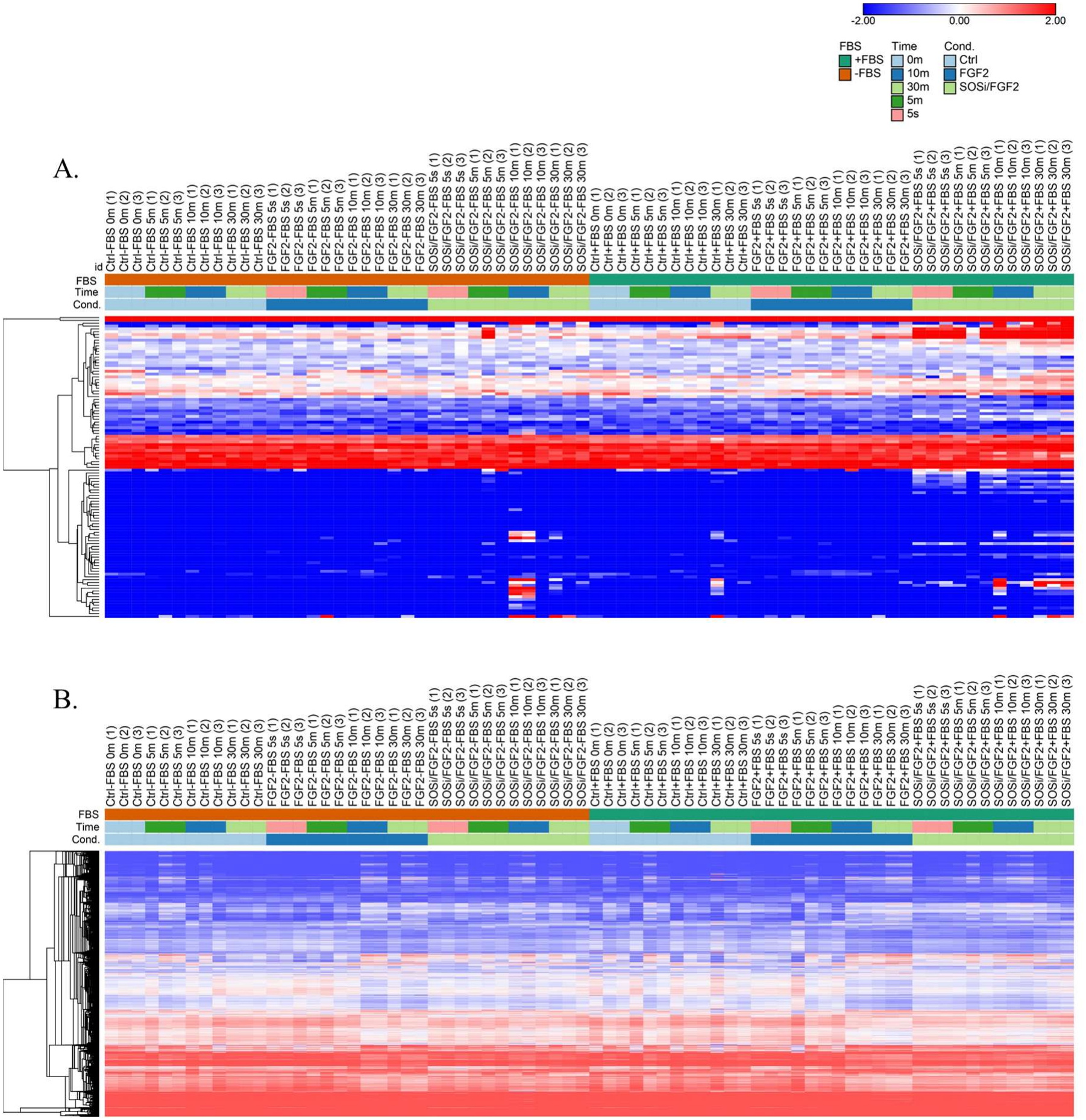
Hierarchical clustering of the proteomes. Limma significant proteome hierarchical clustering (N=111). B: Limma significant phosphoproteome hierarchical clustering (N=10,416).

**Figure 6:**
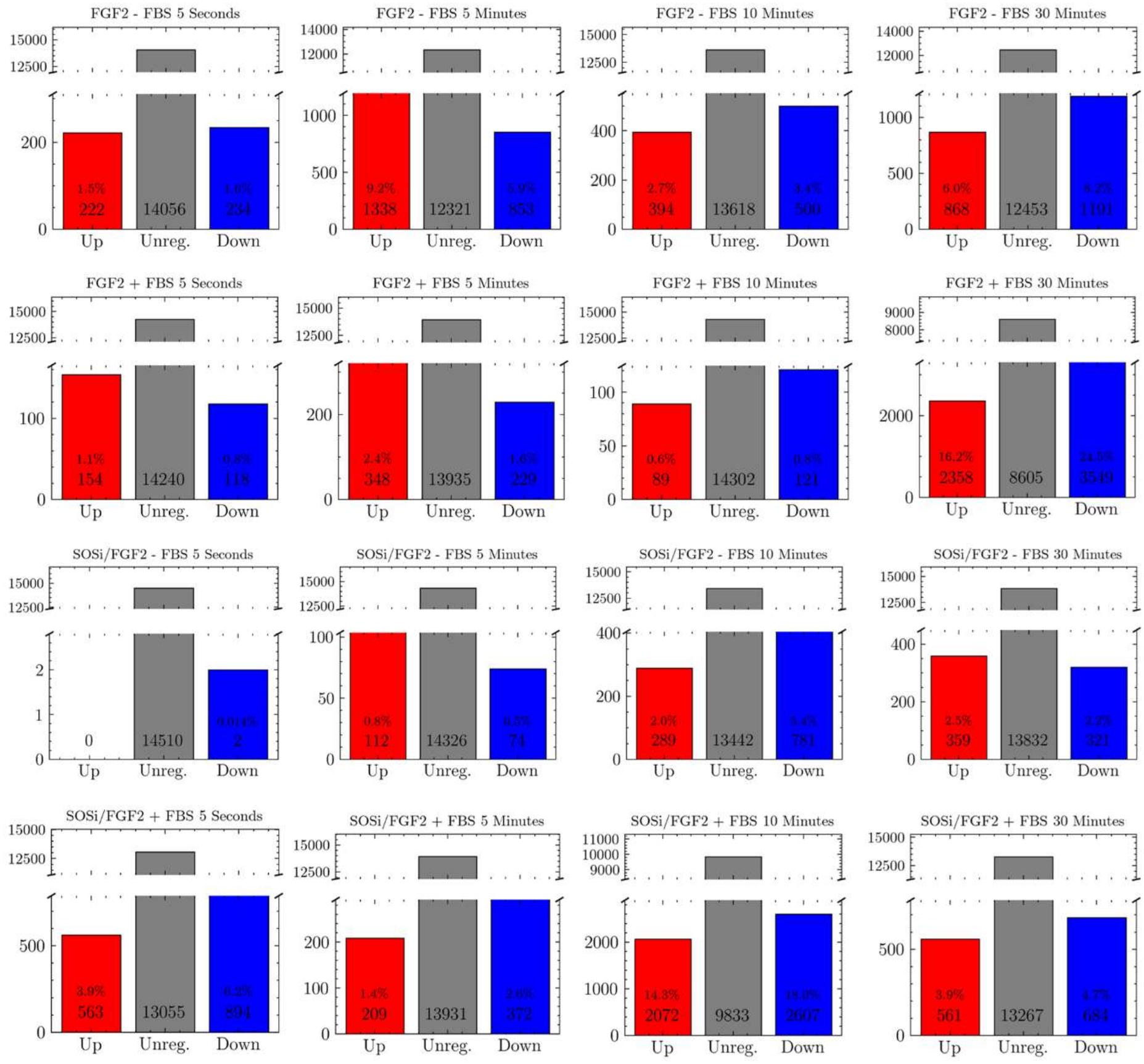
Limma significant phosphoproteome by condition. **Blue bars** represent downregulated phosphosites, **gray bars** indicate phosphosites where no significant change was observed, and the **red bars** represent upregulated phosphosites.

It is also important to note that both the total proteome and the phosphoproteome analyses were derived from the exact same biological samples, split during the sample preparation procedure. The observed stability of the total proteome over the analyzed time span, together with the dynamic changes detected in the phosphoproteome, supports that the phosphorylation changes are biologically meaningful.

### Functional and Pathways Enrichment Analysis

Next, to demonstrate the functional enrichment and the potential pathways involved in this dataset, the Over-representation analysis (ORA) of the proteome dataset was performed using WebGestalt analysis method for Gene Ontology Biological Process (non-redundant) annotations (**Figure 7**). From the 8,786 UniProt Swiss-Prot identifiers provided, after repeated entries were removed, 8,852 were mapped as unique and 7,283 were unambiguously mapped, corresponding to 7,280 unique Entrez Gene identifiers, which formed the basis for the enrichment analysis (**Figure 7A**). Of these, 6,303 genes were annotated to at least one Biological Process term present in the reference background. The reference set comprised 26,281 protein-coding mouse genes, of which 14,490 were annotated to the selected functional categories. Using the defined statistical thresholds, ten Biological Process categories met the significance criteria and are reported (**Figure 7B**).

**Figure 7:**
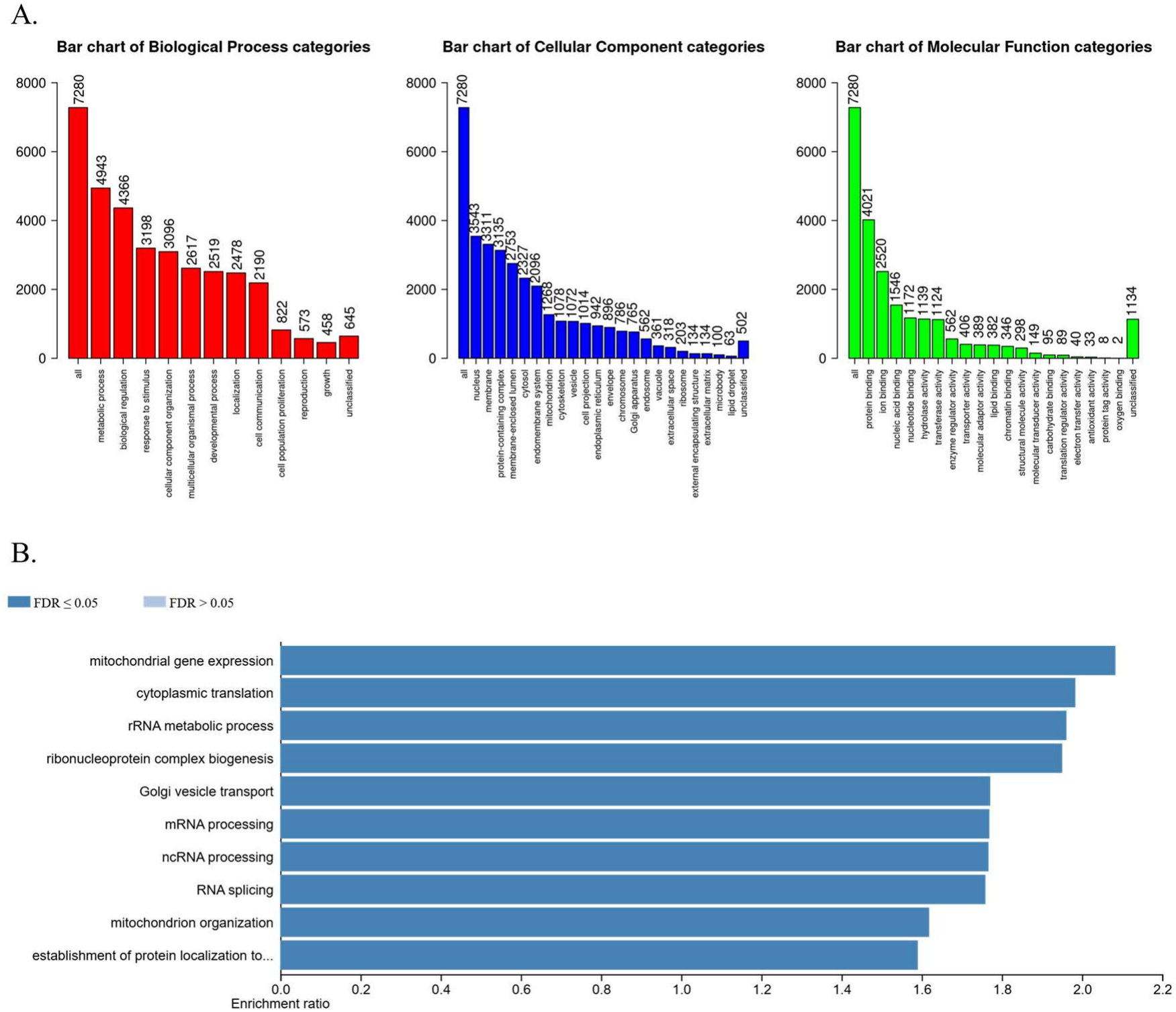
Over-representation analysis (ORA) of the proteome dataset. **A**: The bar chart represents the number of genes fit by three categories. **Red** is the Biological Process category, **Blue** is the Cellular Component category and Green is the Molecular Function category. **B**: Top 10 categories enrichment results. FDR <0.05 is represented in **dark blue**. FDR >0.05 is represented in **light blue**.

Similarly, for the phosphoproteome using a gene-centric representation, the ORA was performed to the phosphosites corresponding genes (**Figure 8**). From 3,986 UniProt Swiss-Prot identifiers, 3,472 were unambiguously mapped, corresponding to 3,471 unique Entrez Gene identifiers used for the analysis (**Figure 8A**). Among these, 2,965 genes were annotated to at least one Gene Ontology Biological Process category present in the reference background. The reference set comprised 26,281 protein-coding mouse genes, of which 14,490 were annotated to the selected functional categories. Ten Biological Process categories met the significance threshold and are reported (**Figure 8B**).

**Figure 8:**
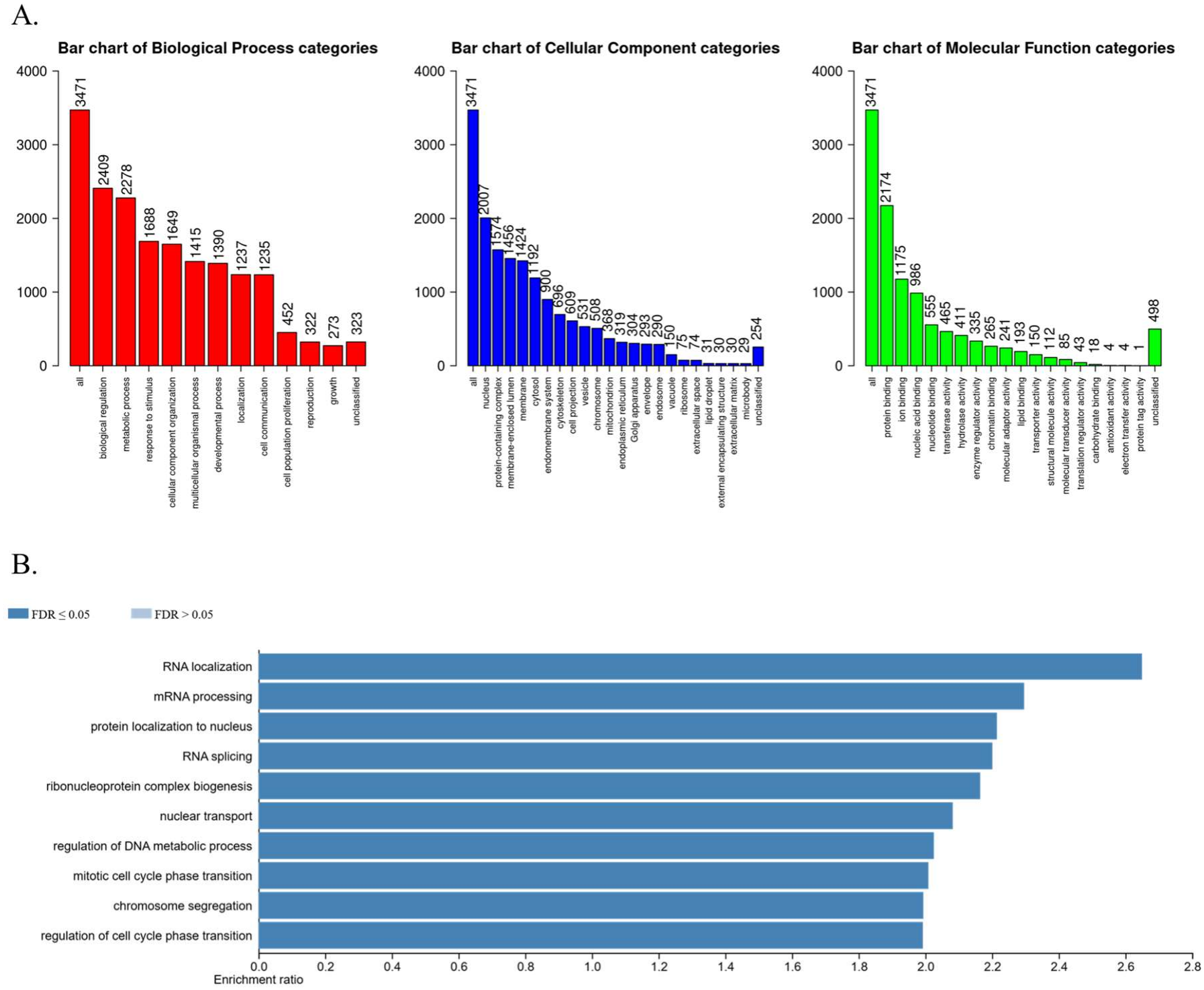
Over-representation analysis (ORA) of the phosphoproteome dataset, in a gene-centric analysis. **A:** The bar chart represents the number of genes fit by three categories. Red is the Biological Process category, Blue is the Cellular Component category and Green is the Molecular Function category. **B:** Top 10 categories enrichment results. FDR <0.05 is represented in **dark blue**. FDR >0.05 is represented **in light blue**.

In sequence, the correspondence of the phosphoproteome phosphosites identified to the kinases can be verified in the **Figure 9**. Mapping of kinases shown in **Figure 9** can be visualized in the **Table 4 (Kinase coverage relative to the known kinome)**.

**Table 4:** Kinase coverage relative to the known kinome. Summary of the kinases identified in this dataset compared with the total number of known kinases, expressed as a percentage of the known kinome. Due to limitations of the Coral kinase annotation tool, *Mus musculus* kinase identifiers were converted to the corresponding Homo sapiens nomenclature to enable compatibility.

| **Kinase family** | **Identified / Total** | **Percentage (%)** |
|---|---|---|
| **AGC** | 53 / 64 | 82.81 |
| **Atypical** | 12 / 30 | 40.00 |
| **CAMK** | 58 / 82 | 70.73 |
| **CK1** | 11 / 12 | 91.67 |
| **CMGC** | 57 / 63 | 90.48 |
| **Other** | 53 / 81 | 65.43 |
| **STE** | 38 / 48 | 79.17 |
| **TLK** | 29 / 43 | 67.44 |
| **TK** | 11 / 94 | 11.70 |
| **RGC** | 0 / 5 | 0.00 |

**Figure 9:**
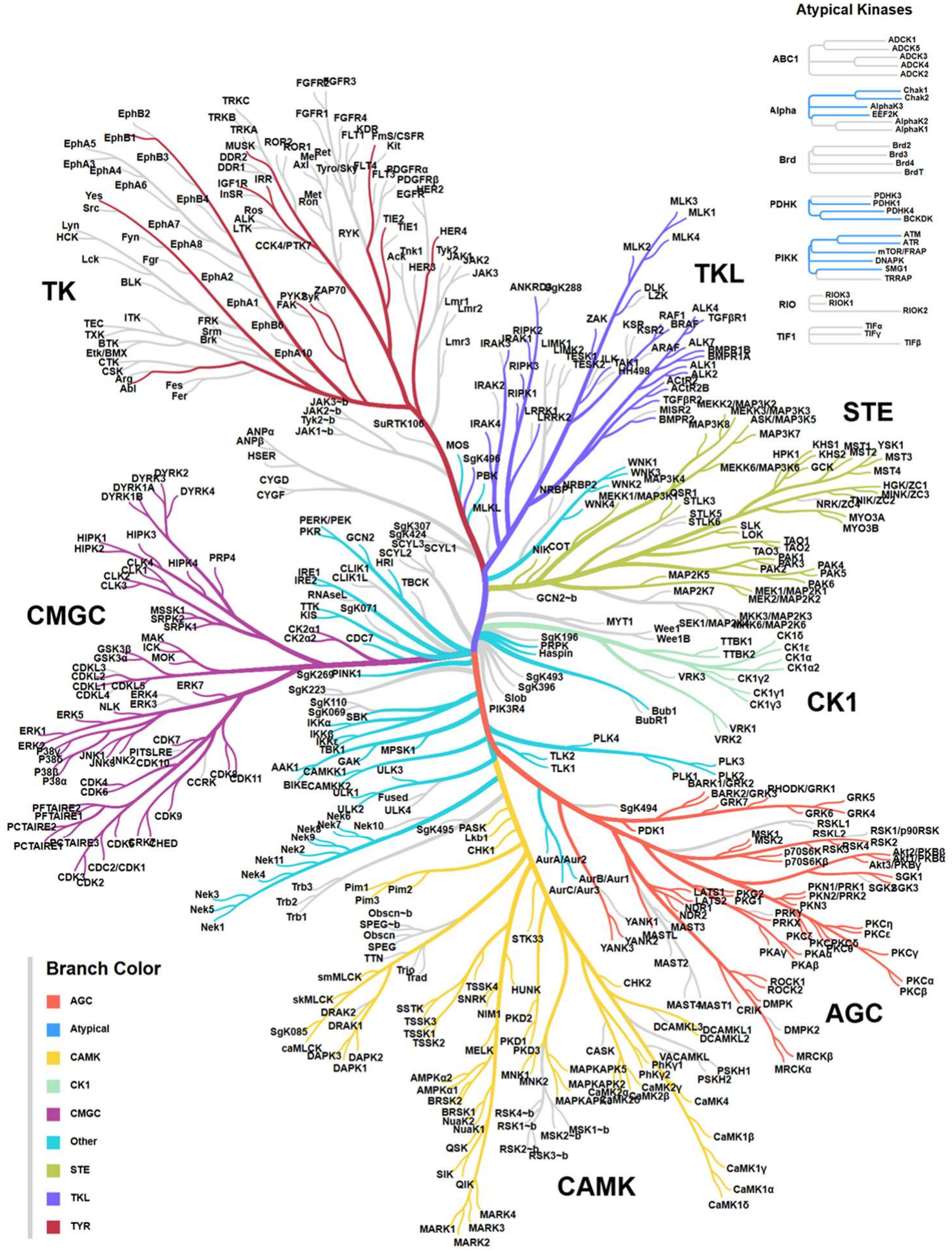
Hierarchical tree of kinases identified in this dataset. Each color represents a kinase branch. **Red**: ACG (PKA/PKG/PKC kinase group), **Blue**: Atypical (Atypical kinases), **Yellow**: CAMK (Calcium/Calmodulin-dependent protein kinases), **Light Green**: CK1 (Casein kinase 1), **Purple**: CMGC (CDK/MAPK/GSK3/CLK kinase group), **Light Blue**: Other (Other kinases), **Olive Green**: STE (Sterile kinases), **Light Purple**: TLK (Tousled-like kinases), **Wine**: TYR (Tyrosine kinases). Names have been converted to HCGM identifiers for *Homo sapiens*.

Finally, gene-centric KEGG pathway enrichment analysis of the kinases identified in the phosphoproteome was performed using STRING, revealed in total 159 pathways enriched, where the top enriched pathway is the MAPK signaling pathway (FDR 1.76 × 10^−41^) and the top 20 are shown in the **Table 5 (Top 20 KEGG pathway enrichment analysis)**.

**Table 5:** Top 20 KEGG pathway enrichment analysis of kinases identified in the phosphoproteome. The table summarizes the top 20 enriched KEGG pathways among all kinases identified in the phosphoproteomic dataset, based on enrichment analysis performed using STRING.

| #term ID | term description | observed gene count | background gene count | strength | signal | false discovery rate |
|---|---|---|---|---|---|---|
| mmu04010 | MAPK signaling pathway | 54 | 287 | 121 | 525 | 1.76E-41 |
| mmu04722 | Neurotrophin signaling pathway | 32 | 114 | 139 | 514 | 7.46E-28 |
| mmu04068 | FoxO signaling pathway | 30 | 129 | 131 | 438 | 5.42E-24 |
| mmu04660 | T cell receptor signaling pathway | 28 | 103 | 137 | 463 | 6.44E-24 |
| mmu04914 | Progesterone-mediated oocyte maturation | 26 | 90 | 14 | 456 | 1.13E-22 |
| mmu04012 | ErbB signaling pathway | 25 | 81 | 143 | 462 | 2.27E-22 |
| mmu05170 | Human immunodeficiency virus 1 infection | 33 | 224 | 111 | 34 | 1.89E-21 |
| mmu05135 | Yersinia infection | 26 | 122 | 127 | 38 | 5.86E-20 |
| mmu04218 | Cellular senescence | 28 | 168 | 116 | 338 | 3.55E-19 |
| mmu04140 | Autophagy - animal | 26 | 136 | 122 | 355 | 5.29E-19 |
| mmu04910 | Insulin signaling pathway | 25 | 133 | 121 | 342 | 4.65E-18 |
| mmu05161 | Hepatitis B | 26 | 160 | 115 | 318 | 1.68E-17 |
| mmu05235 | PD-L1 expression and PD-1 checkpoint pathway in cancer | 21 | 86 | 133 | 358 | 6.68E-17 |
| mmu01522 | Endocrine resistance | 21 | 91 | 13 | 346 | 1.7E-16 |
| mmu04668 | TNF signaling pathway | 22 | 112 | 123 | 324 | 4.22E-16 |
| mmu04912 | GnRH signaling pathway | 20 | 85 | 131 | 339 | 8.12E-16 |
| mmu05163 | Human cytomegalovirus infection | 28 | 237 | 101 | 263 | 8.12E-16 |
| mmu05212 | Pancreatic cancer | 19 | 72 | 136 | 349 | 9.21E-16 |
| mmu04150 | mTOR signaling pathway | 24 | 156 | 113 | 292 | 1.12E-15 |
| mmu05200 | Pathways in cancer | 38 | 527 | 8 | 203 | 1.51E-15 |

Taken altogether, this dataset provides a foundation for understanding Y1 signaling and the dynamics of this cell in response to important stimuli such as FGF2 and GEF SOS inhibition. This data can be applied to large-scale functional analyses, the discovery of new therapeutic targets that can be translated to similar cell lines, and also to integrate dynamic models, such as ordinary differential equation models.

## Data Records

1. The raw files from the mass spectrometry analysis will be available only in the final version of this manuscript.
2. The tables containing the matrixes will be stored as supplementary data and will be available only in the final version of this manuscript.

## Code Availability

For this version of the manuscript the code used in the is not available for the preprint version.

## Acknowledgements

We thank Hugo A. Armelin for insightful discussions and valuable intellectual contributions throughout the development of this work. We also gratefully acknowledge grant/ support from the São Paulo Research Foundation (FAPESP; Grants #13/07467-1, #20/10329-3, #19/21619-5, #20/08555-5, #19/24580-2, and #21/04355-4), the Coordination for the Improvement of Higher Education Personnel (CAPES; Finance Code 001), NIBN (National Institute of Biomedical Innovation, Health and Nutrition) and the Butantan Institute.

