## Supplementary Information for "A comprehensive temporal proteome and phosphoproteome dataset of Y1 adrenocortical cells under FGF2 stimulation and SOS inhibition"

    - [1] Center of Toxins, Immune-Response and Cell Signaling (CeTICS), Butantan Institute, São Paulo, 05503-900, Brazil.
    - [2] Cell Cycle Laboratory, Butantan Institute, São Paulo, 05503-900, Brazil.[3] Bioinformatics Graduate Program, University of São Paulo, Brazil
    - [3] Laboratory of Artificial Intelligence and Inference in Complex Data (Recod.ai), Institute of Computing, University of Campinas, Campinas, 13083-852, Brazil.
    - [4] AI Center for Health and Biomedical Research (ArCHER), National Institutes of Biomedical Innovation, Health and Nutrition, Osaka, 566-0002, Japan.
    - [5] Laboratory of Proteomics for Drug Discovery, Center for Drug Design Research (CDDR), National Institutes of Biomedical Innovation, Health and Nutrition, Ibaraki, 567-0085, Japan.
    - [‡] Current affiliation: Laboratory of Computational Biomedicine, Graduate School of Science, Technology and Innovation, Kobe University, Kobe, 650-0017, Japan.
    - [§] Current affiliation: Department of Mechanical and System Engineering, Graduate School of Engineering, University of Fukui, Fukui, 910-8507, Japan.
    - * Correspondence Author
    - † These authors jointly supervised this work



- **Keywords**: Phosphoproteomics, Y1 Adrenocortical Cell, Cancer Research, Cell Signaling, Mass Spectrometry

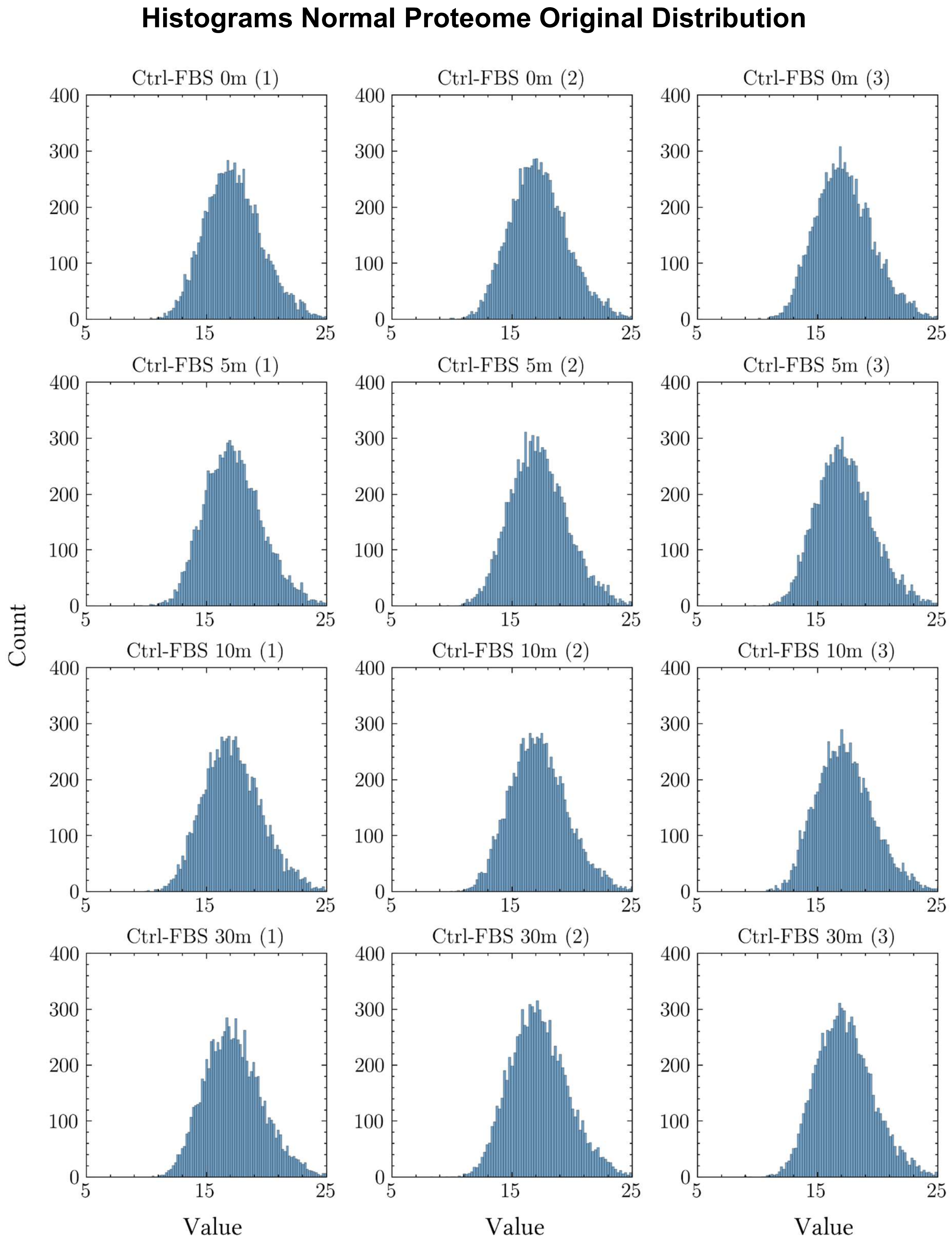


**Figure S1:** Histogram analysis of the frequency distribution of normal proteome data Control minus FBS.

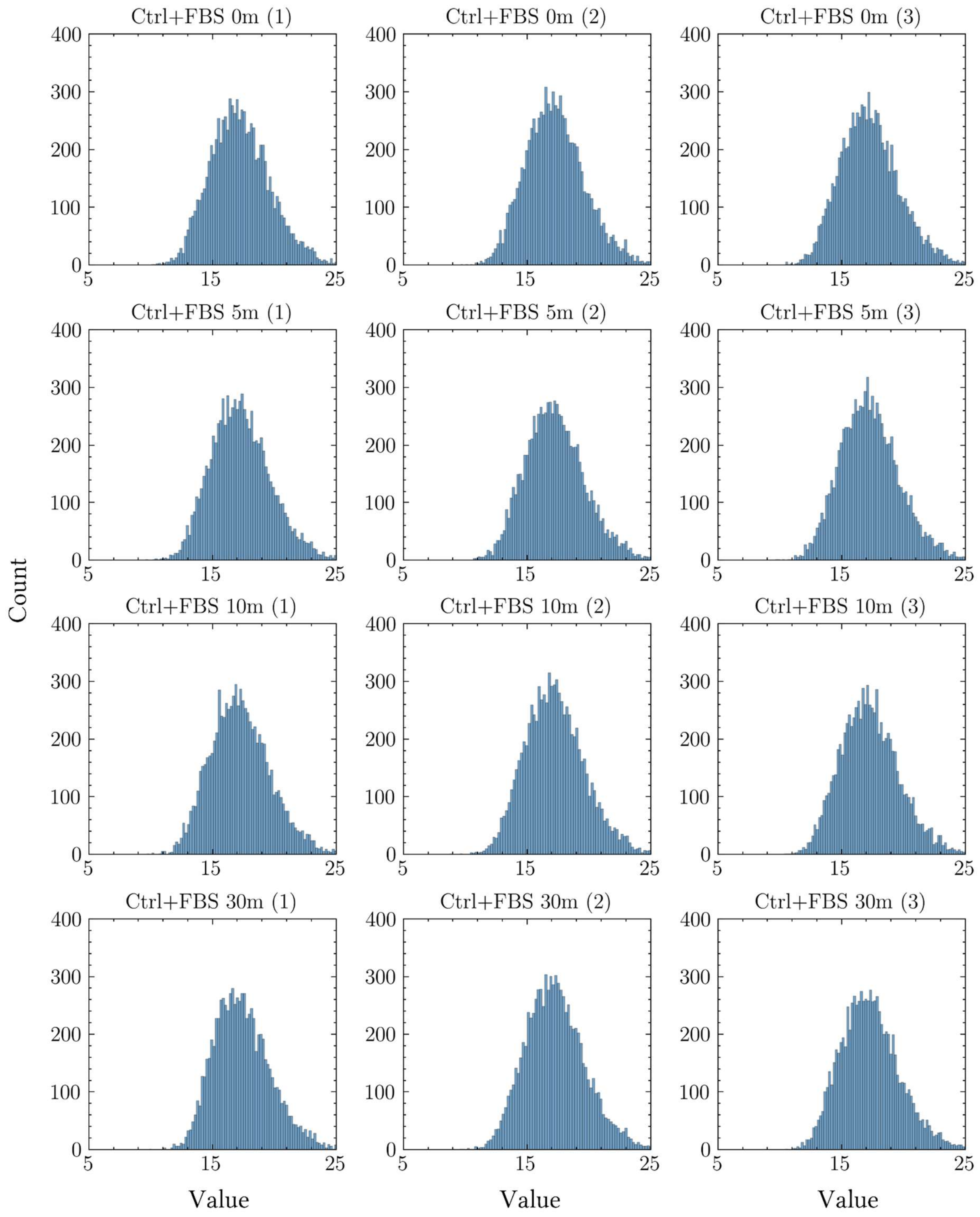


**Figure S2:** Histogram analysis of the frequency distribution of normal proteome data Control plus FBS.

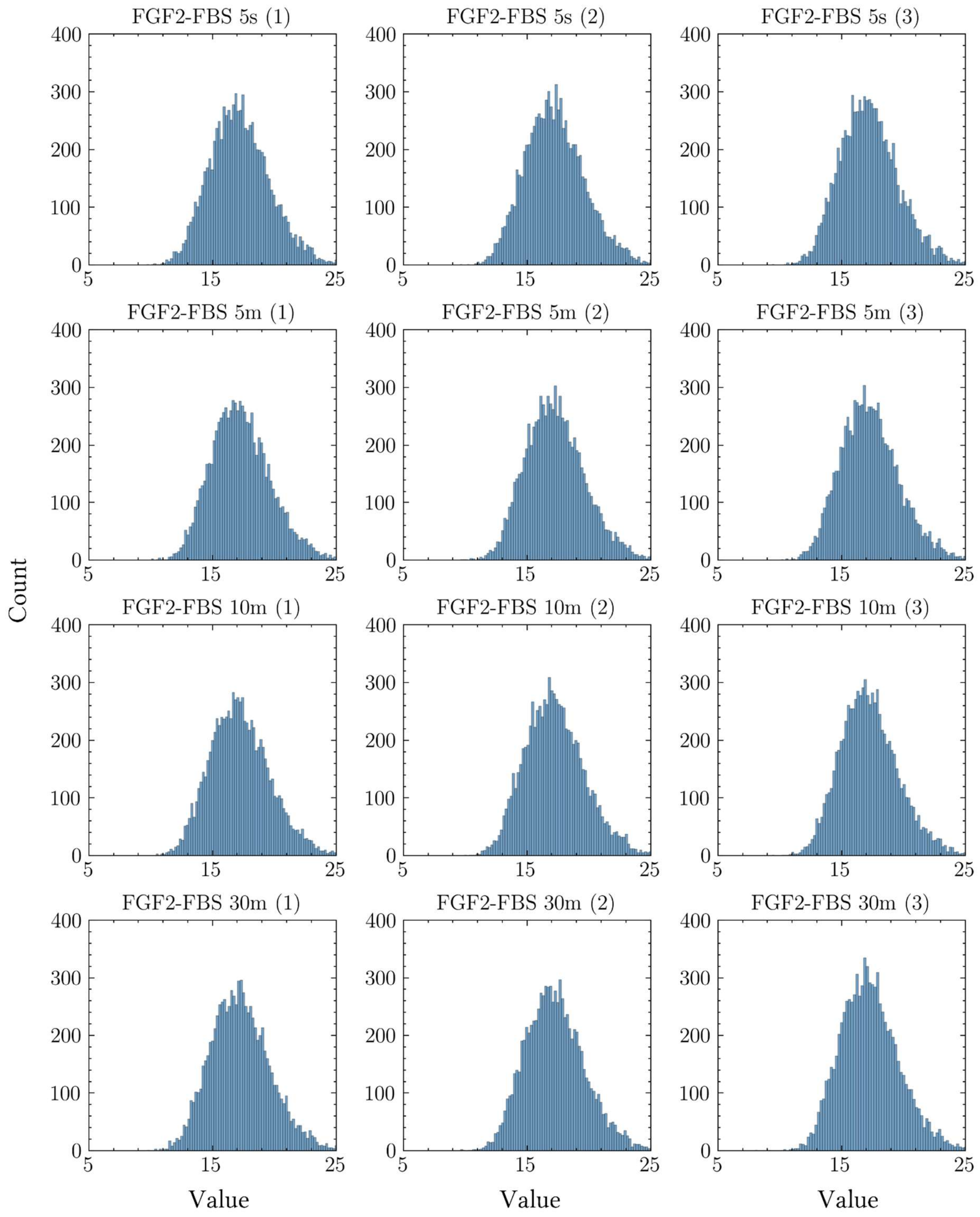


**Figure S3:** Histogram analysis of the frequency distribution of normal proteome data FGF2 minus FBS.

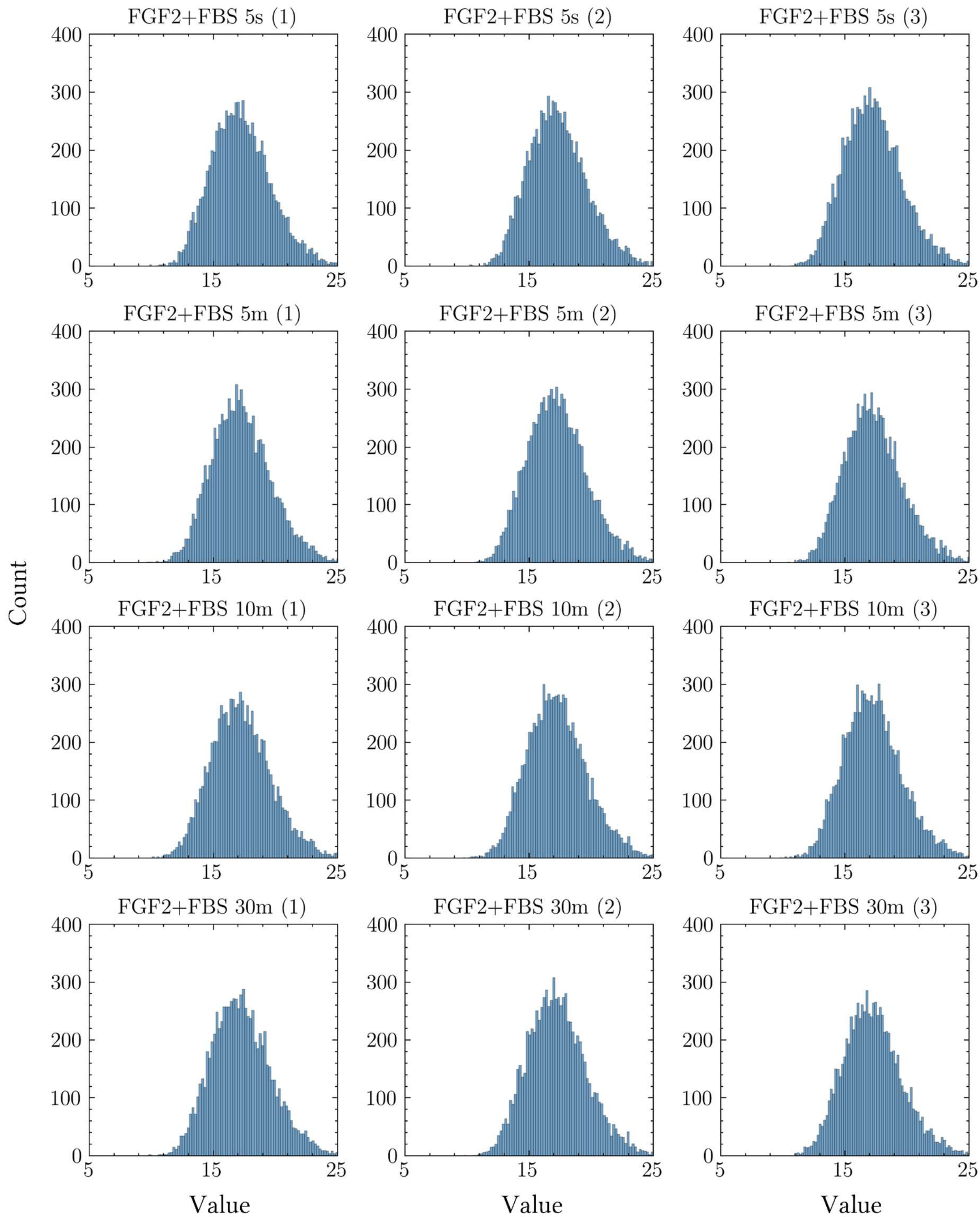


**Figure S4:** Histogram analysis of the frequency distribution of normal proteome data FGF2 plus FBS.

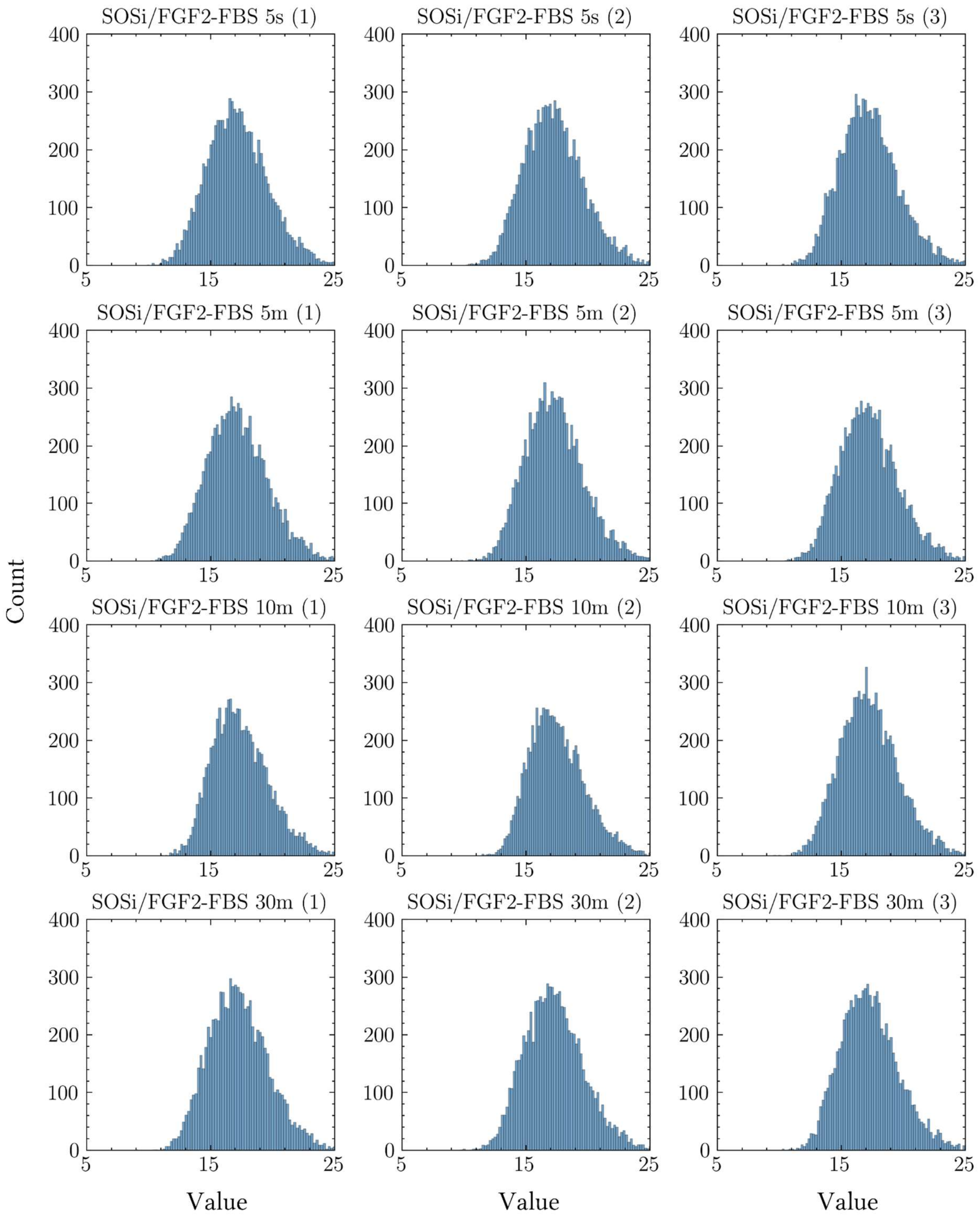


**Figure S5:** Histogram analysis of the frequency distribution of normal proteome data SOSi/FGF2 minus FBS.

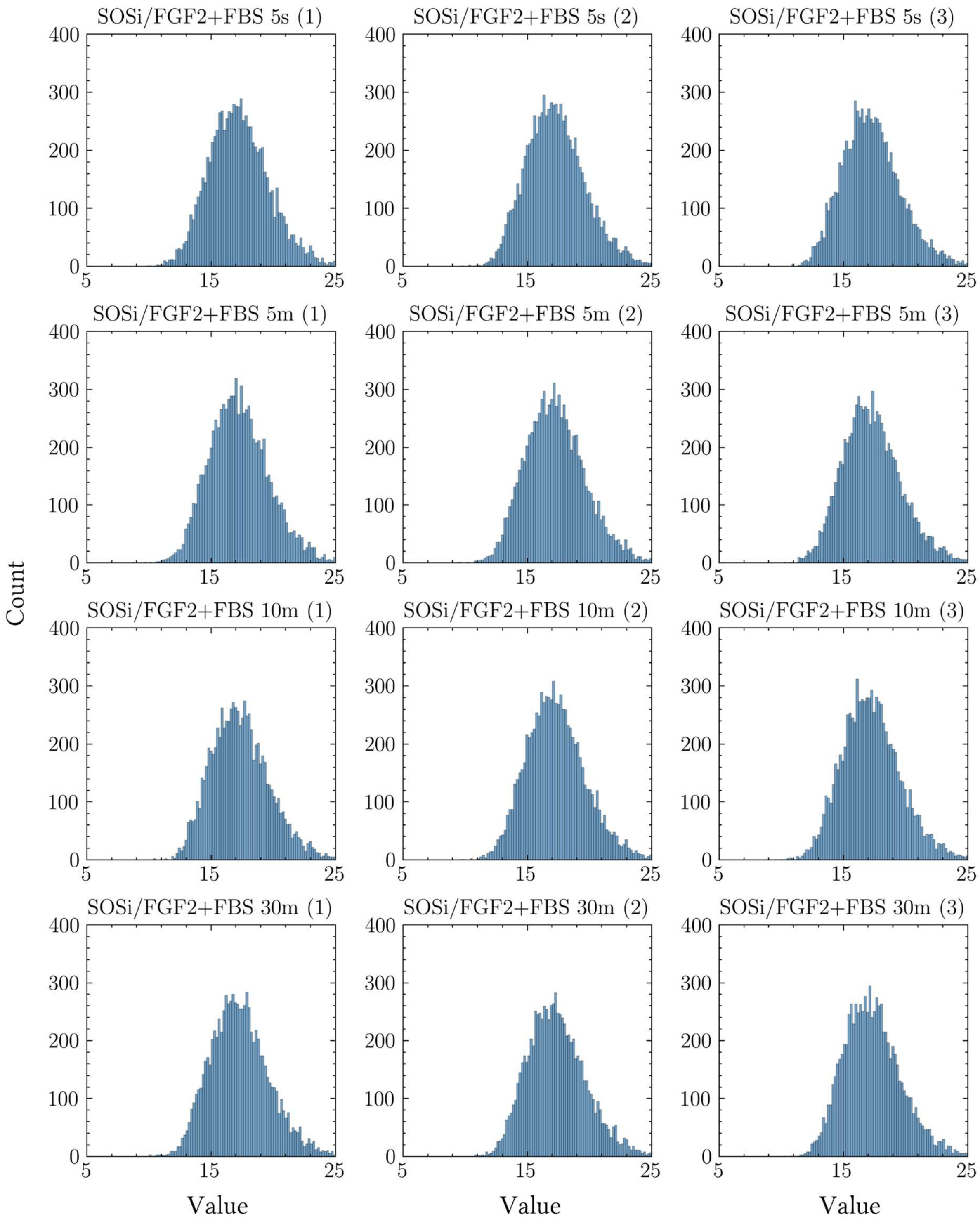


**Figure S6:** Histogram analysis of the frequency distribution of normal proteome data SOSi/FGF2 plus FBS.

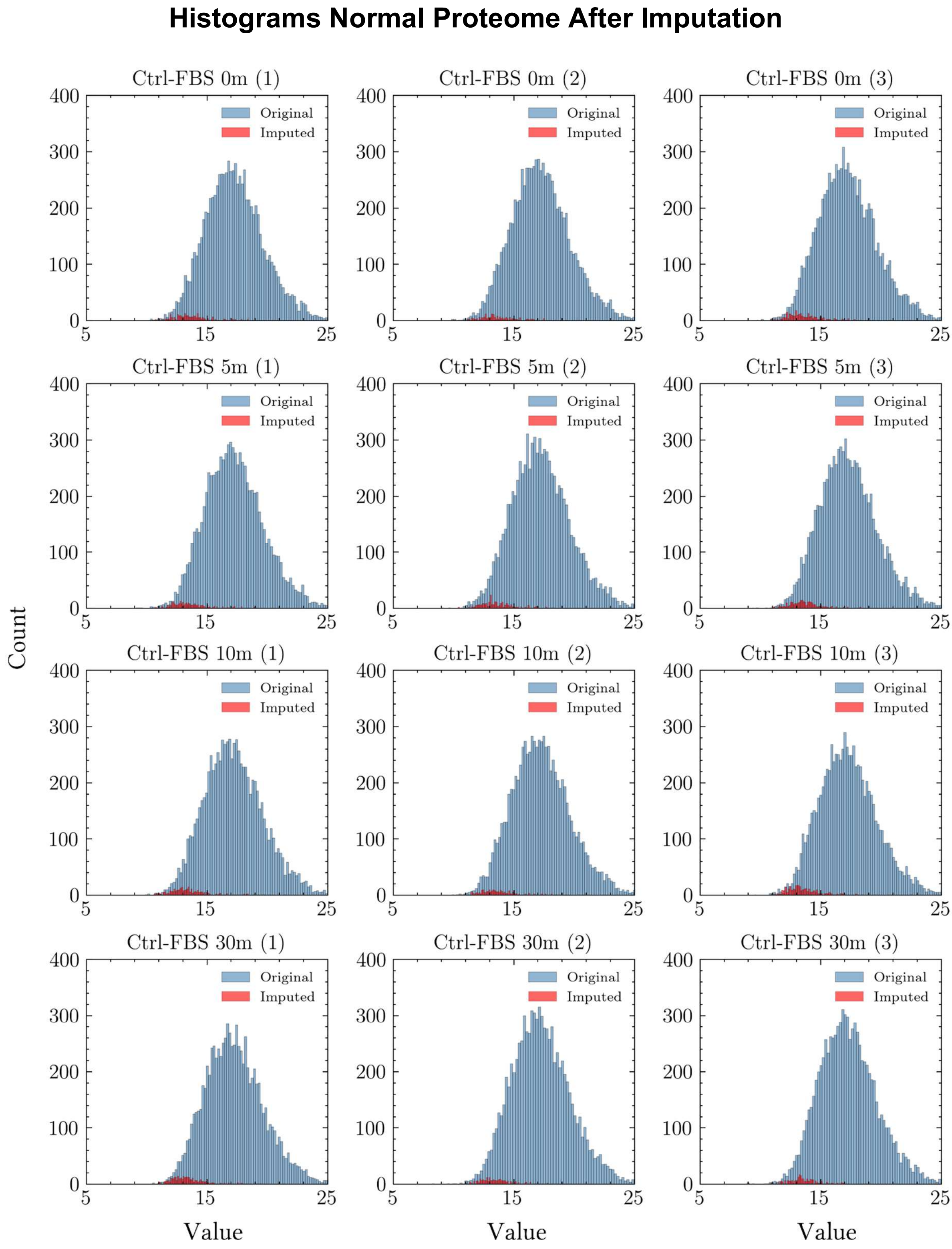


**Figure S7:** Histogram analysis of the frequency distribution of BPCA imputed normal proteome data Control minus FBS.

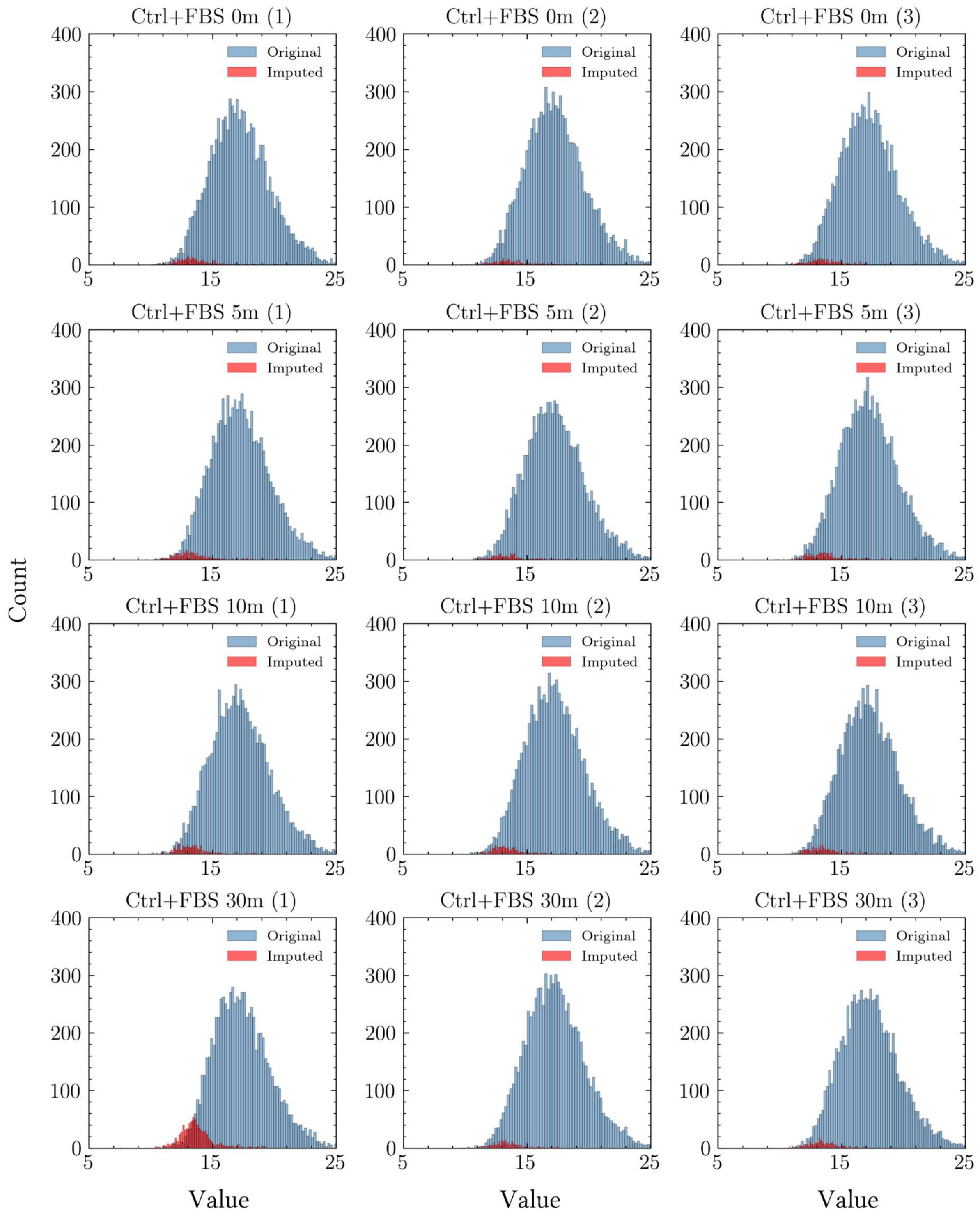


**Figure S8:** Histogram analysis of the frequency distribution of BPCA imputed normal proteome data Control plus FBS.

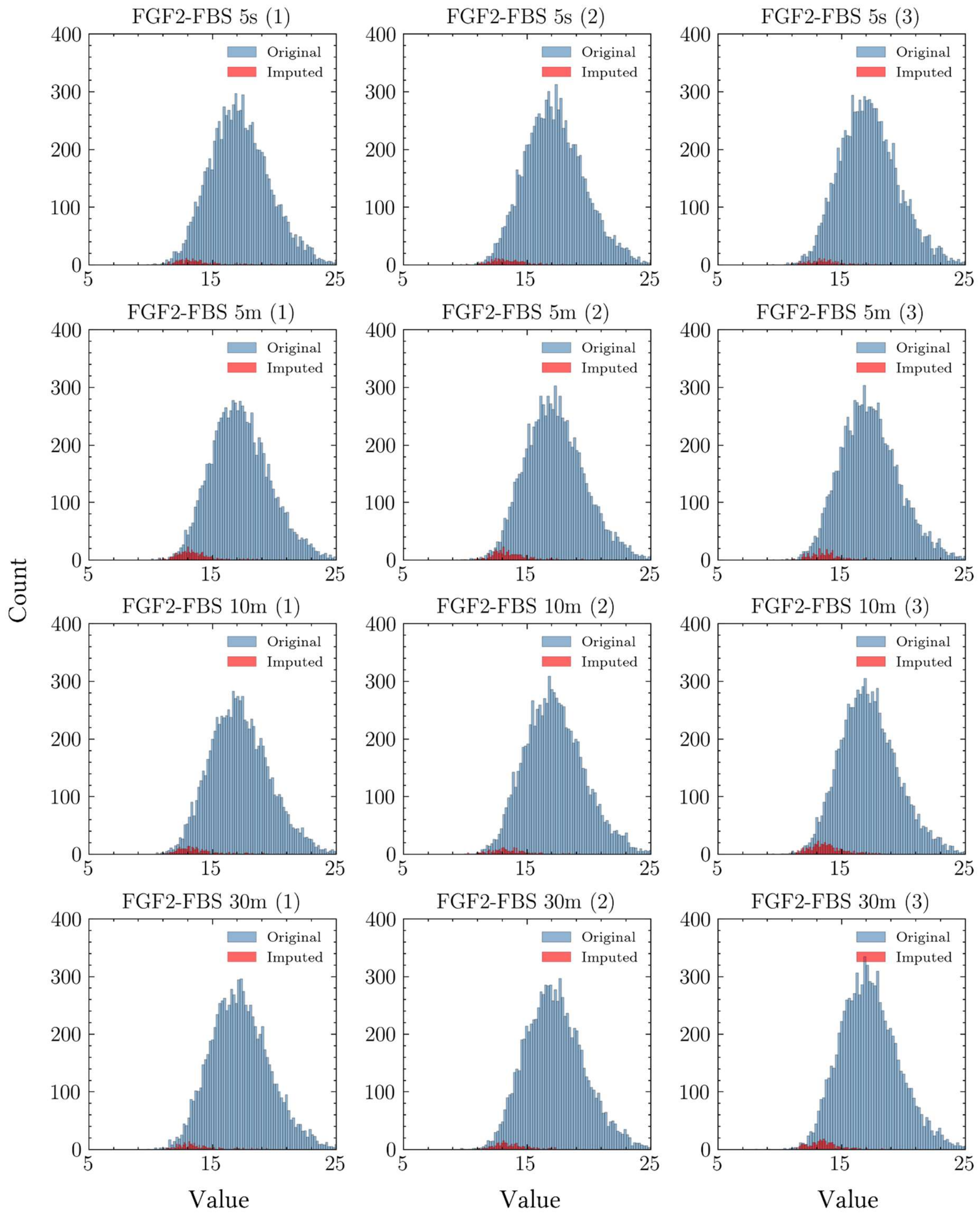


**Figure S9:** Histogram analysis of the frequency distribution of BPCA imputed normal proteome data FGF2 minus FBS.

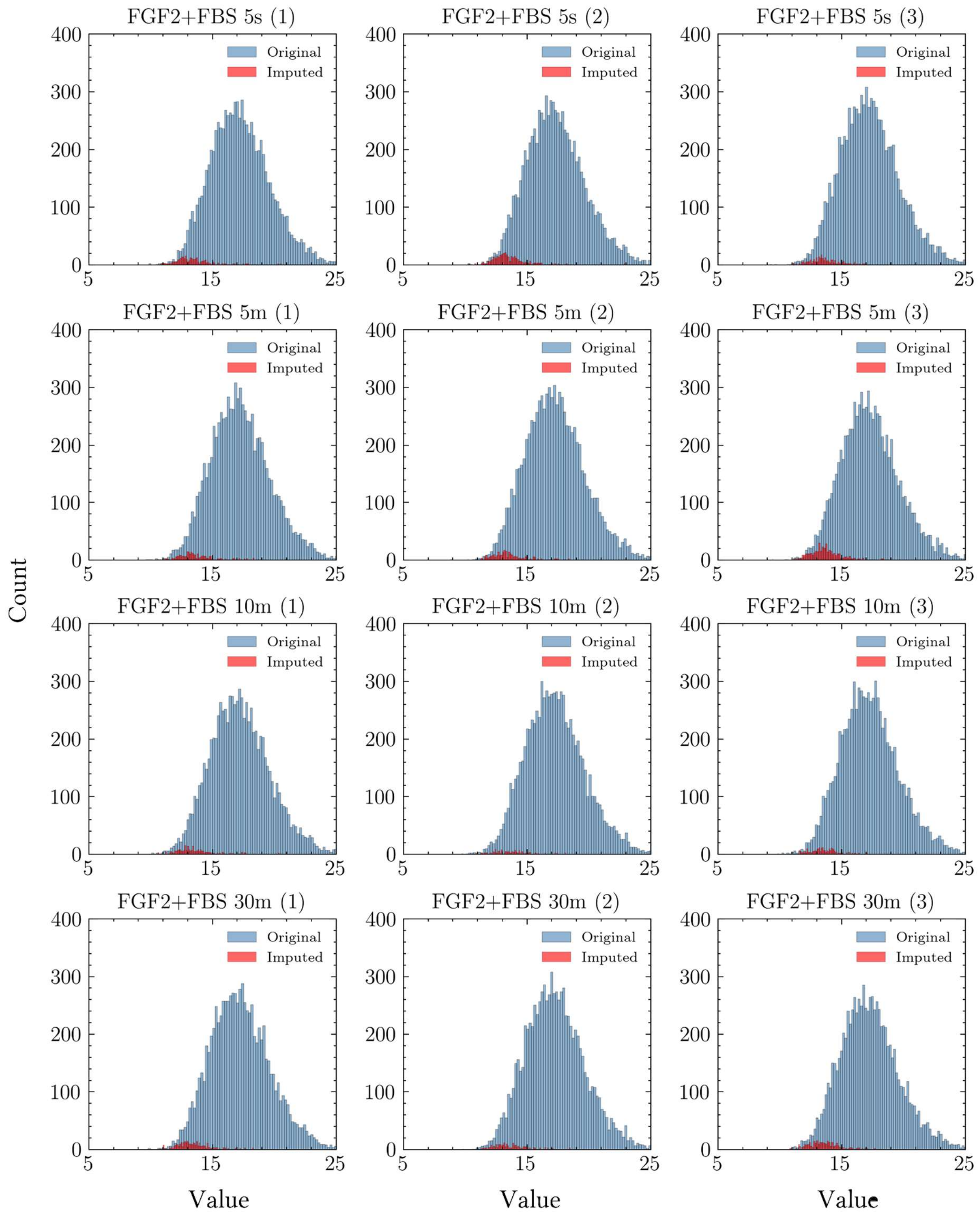


**Figure S10:** Histogram analysis of the frequency distribution of BPCA imputed normal proteome data FGF2 plus FBS.

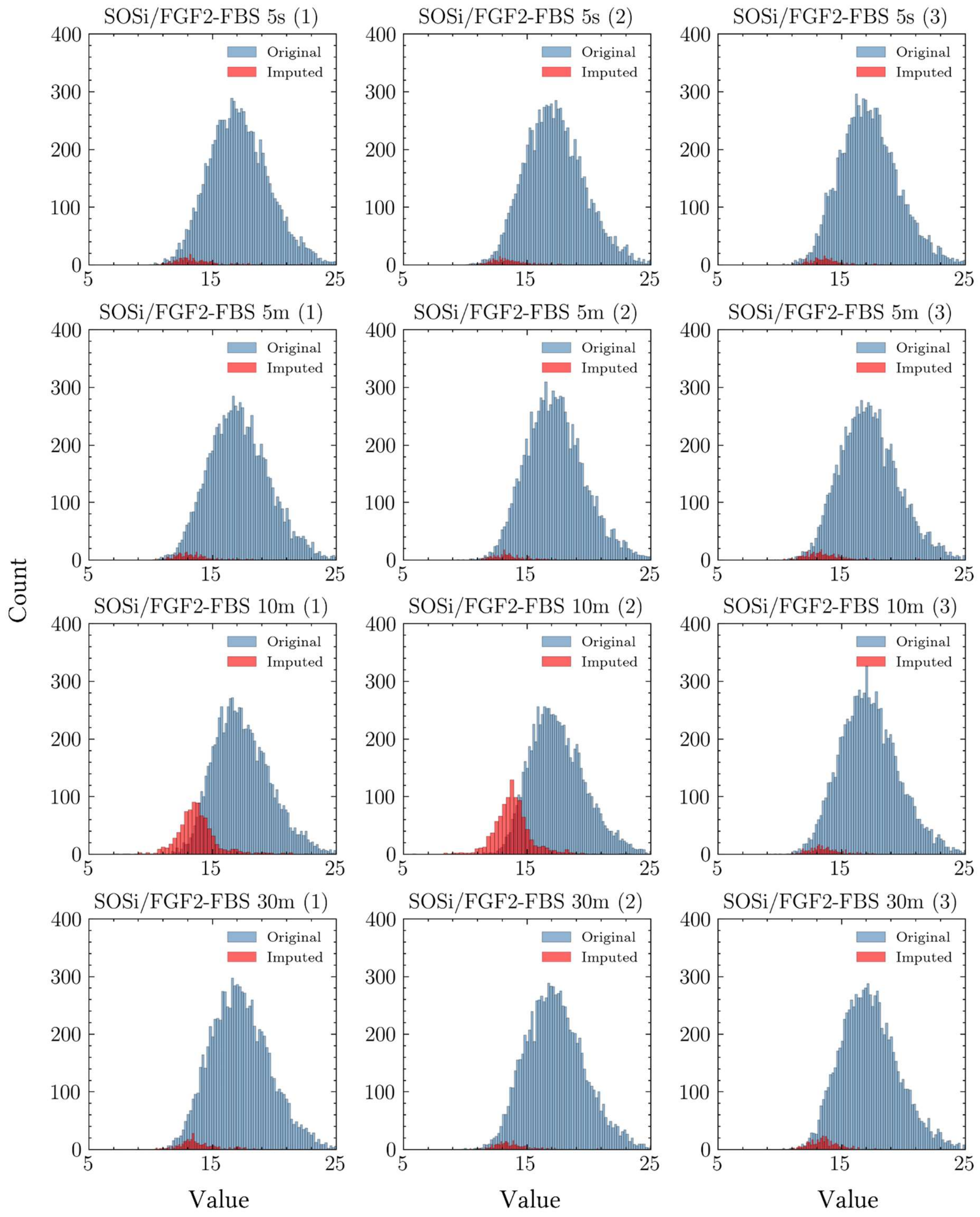


**Figure S11:** Histogram analysis of the frequency distribution of BPCA imputed normal proteome data SOSi/FGF2 minus FBS.

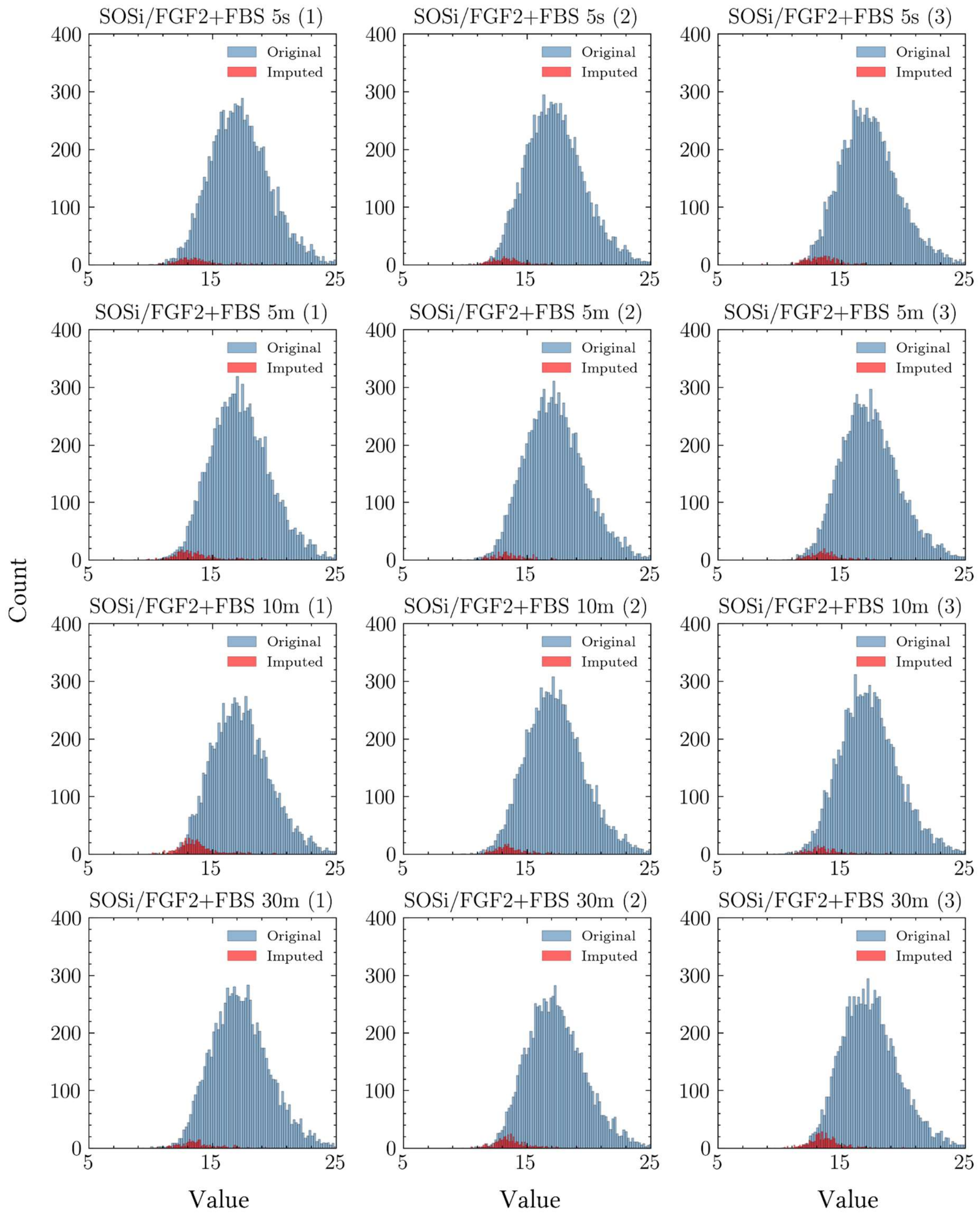


**Figure S12:** Histogram analysis of the frequency distribution of BPCA imputed normal proteome data SOSi/FGF2 plus FBS.

### Histograms Phosphoproteome Original Distribution

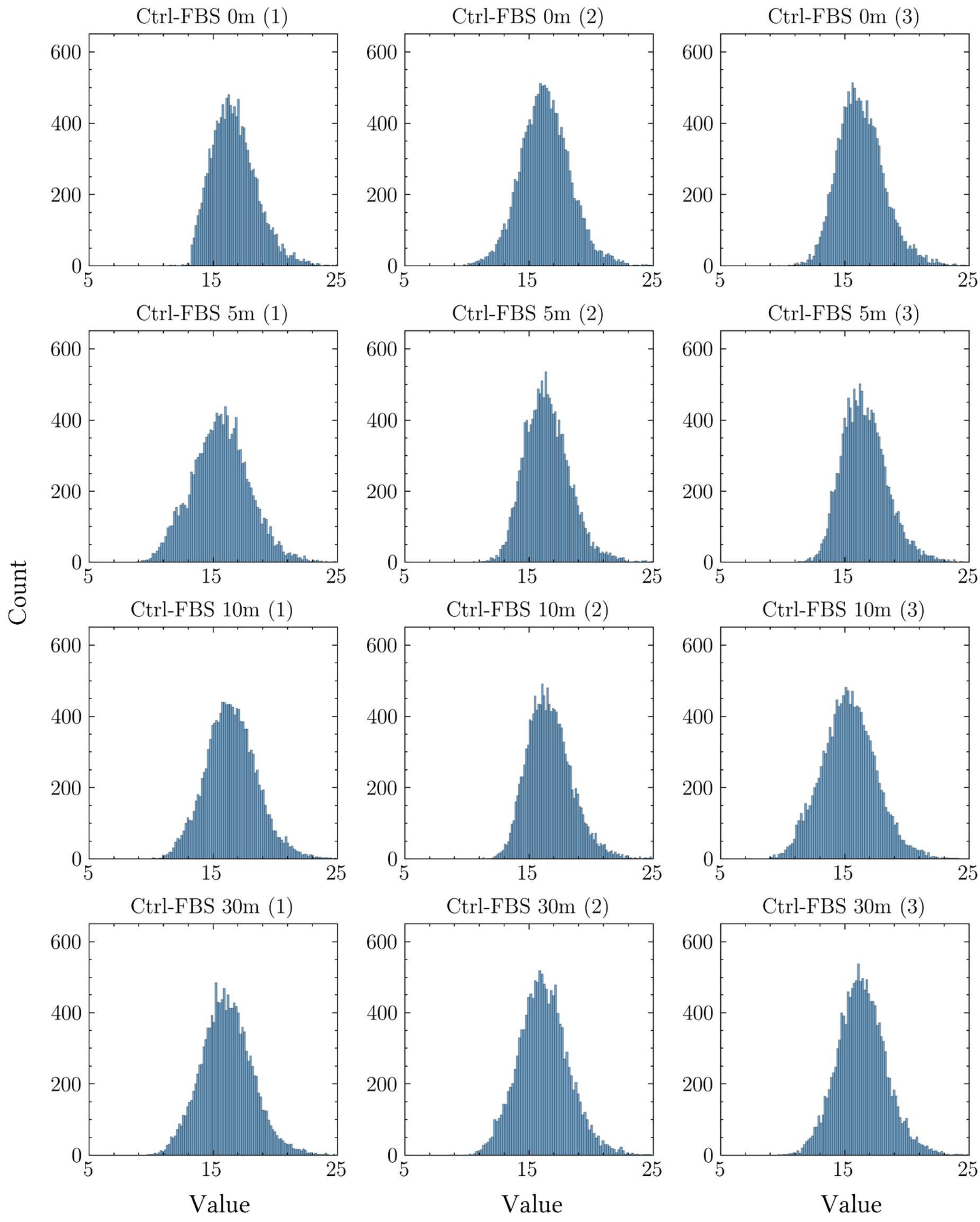


**Figure S13:** Histogram analysis of the frequency distribution of phosphoproteome data Control minus FBS.

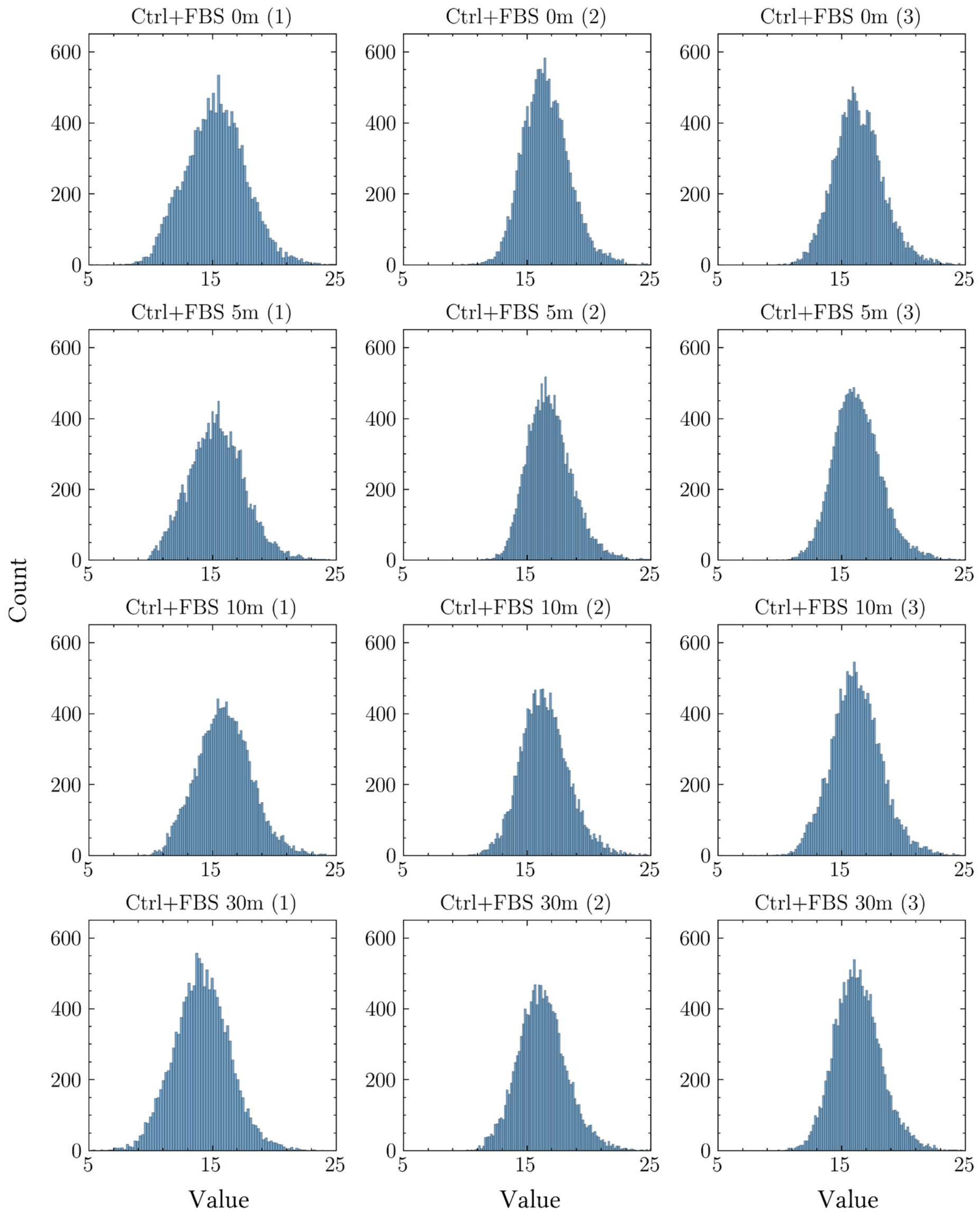


**Figure S14:** Histogram analysis of the frequency distribution of phosphoproteome data Control plus FBS.

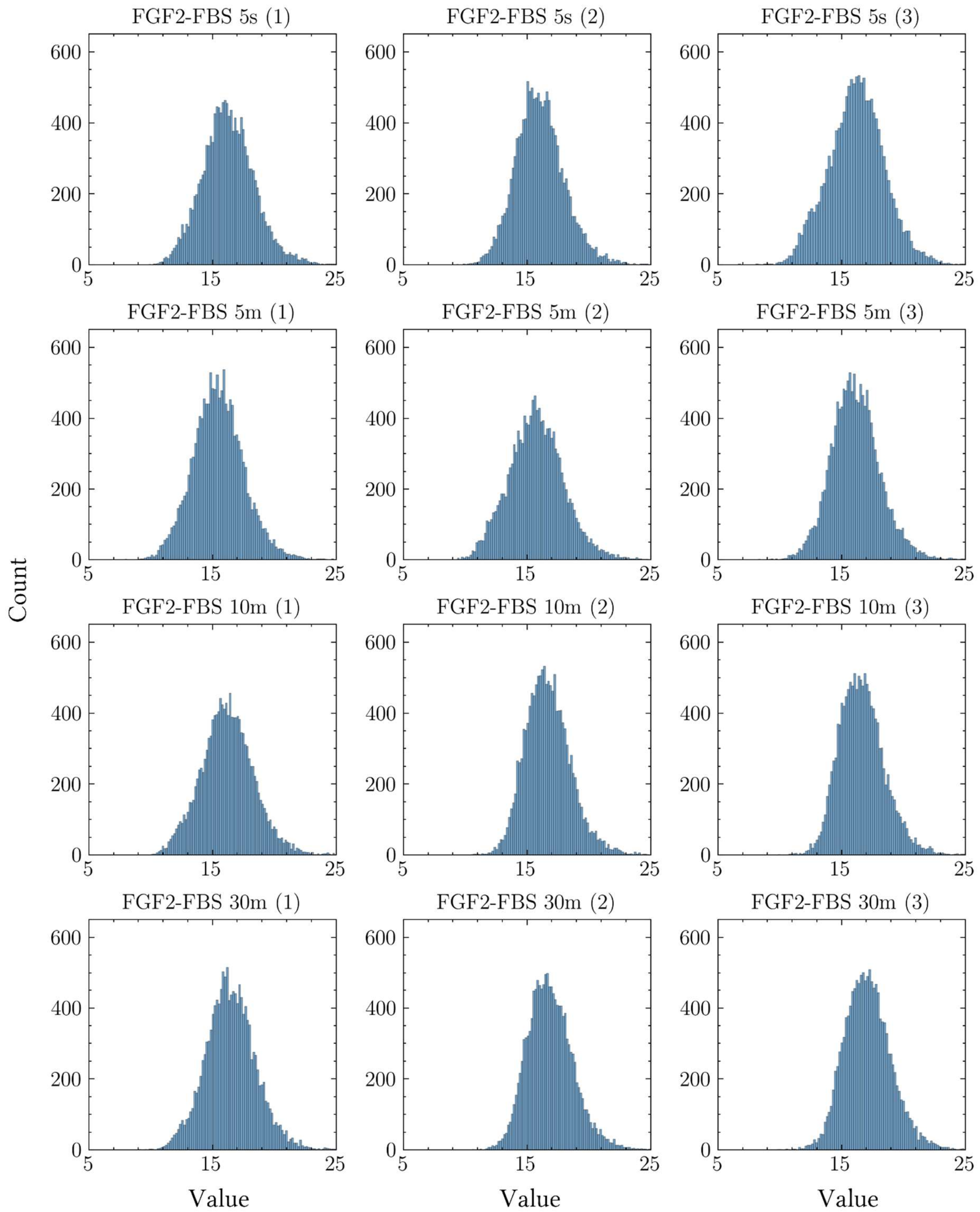


**Figure S15:** Histogram analysis of the frequency distribution of phosphoproteome data FGF2 minus FBS.

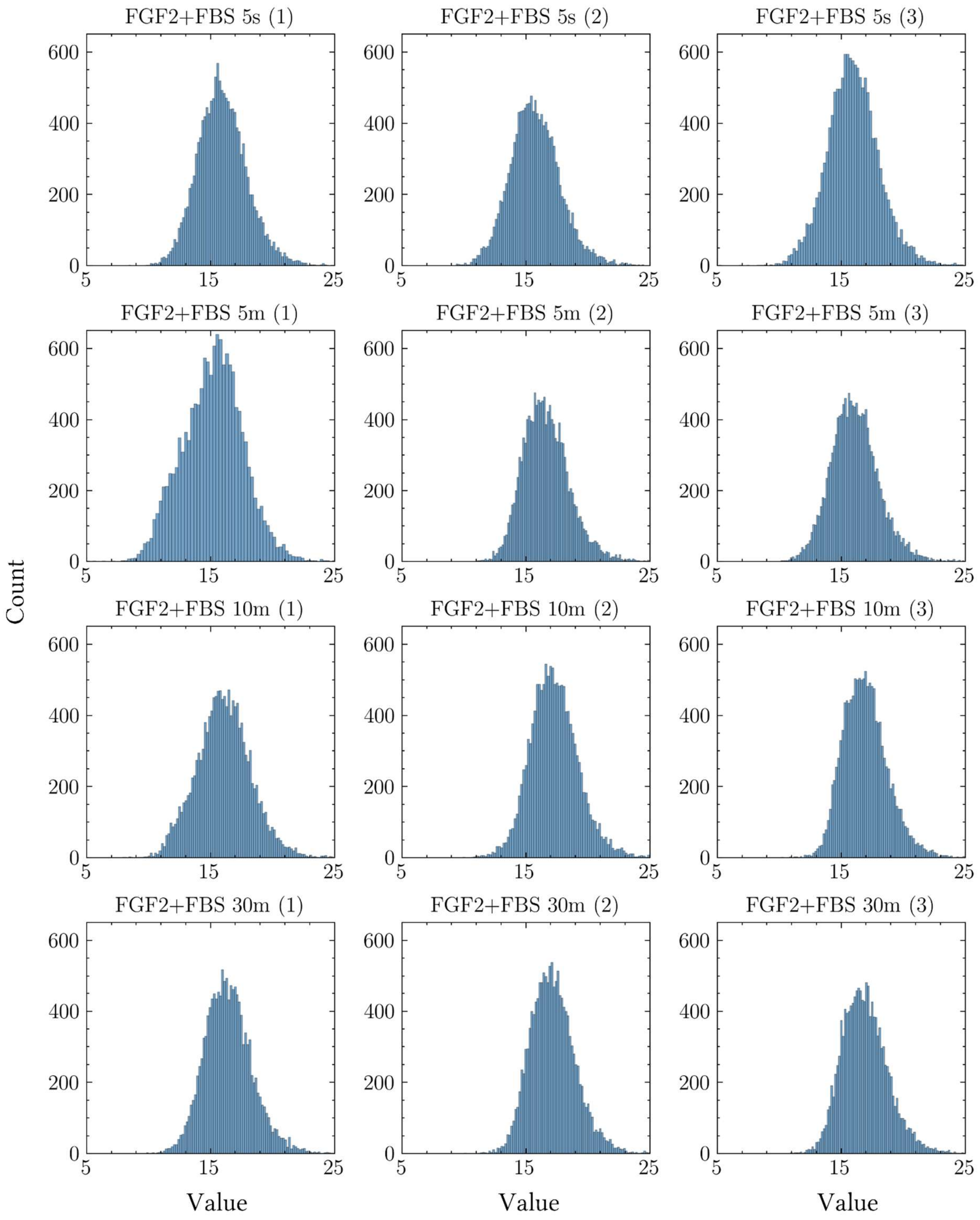


**Figure S16:** Histogram analysis of the frequency distribution of phosphoproteome data FGF2 plus FBS.

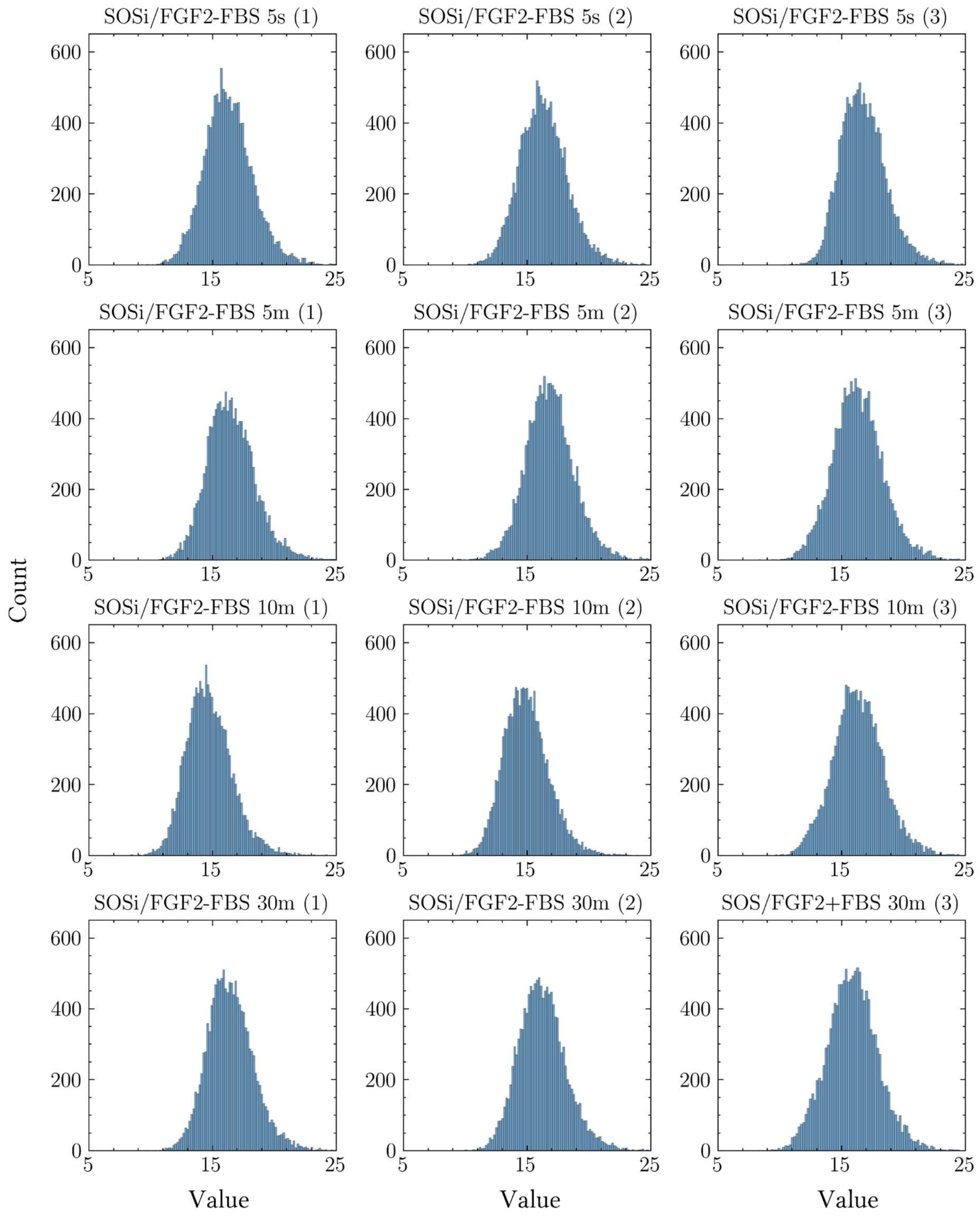


**Figure S17:** Histogram analysis of the frequency distribution of phosphoproteome data SOSi/FGF2 minus FBS.

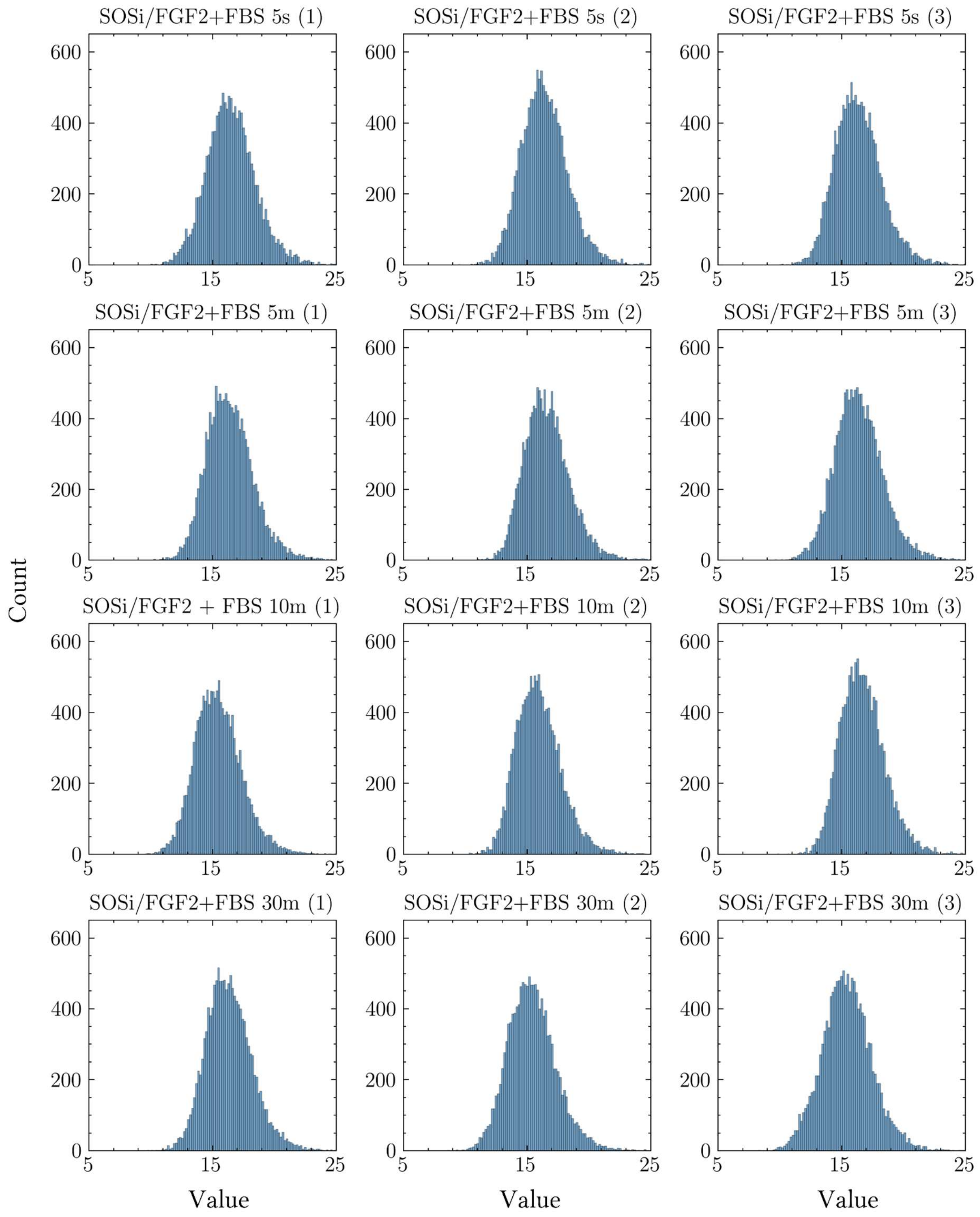


**Figure S18:** Histogram analysis of the frequency distribution of phosphoproteome data SOSi/FGF2 plus FBS.

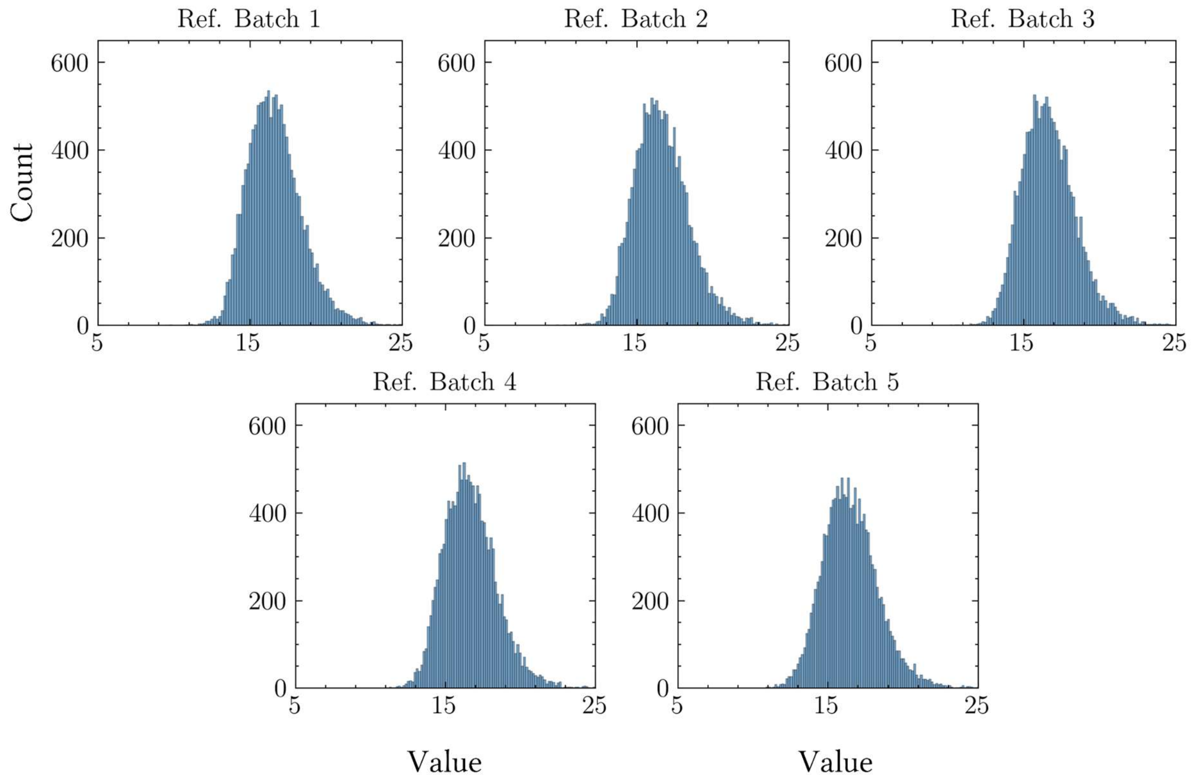


**Figure S19:** Histogram analysis of the frequency distribution of phosphoproteome data Reference Samples..

### Histograms Phosphoproteome After Imputation

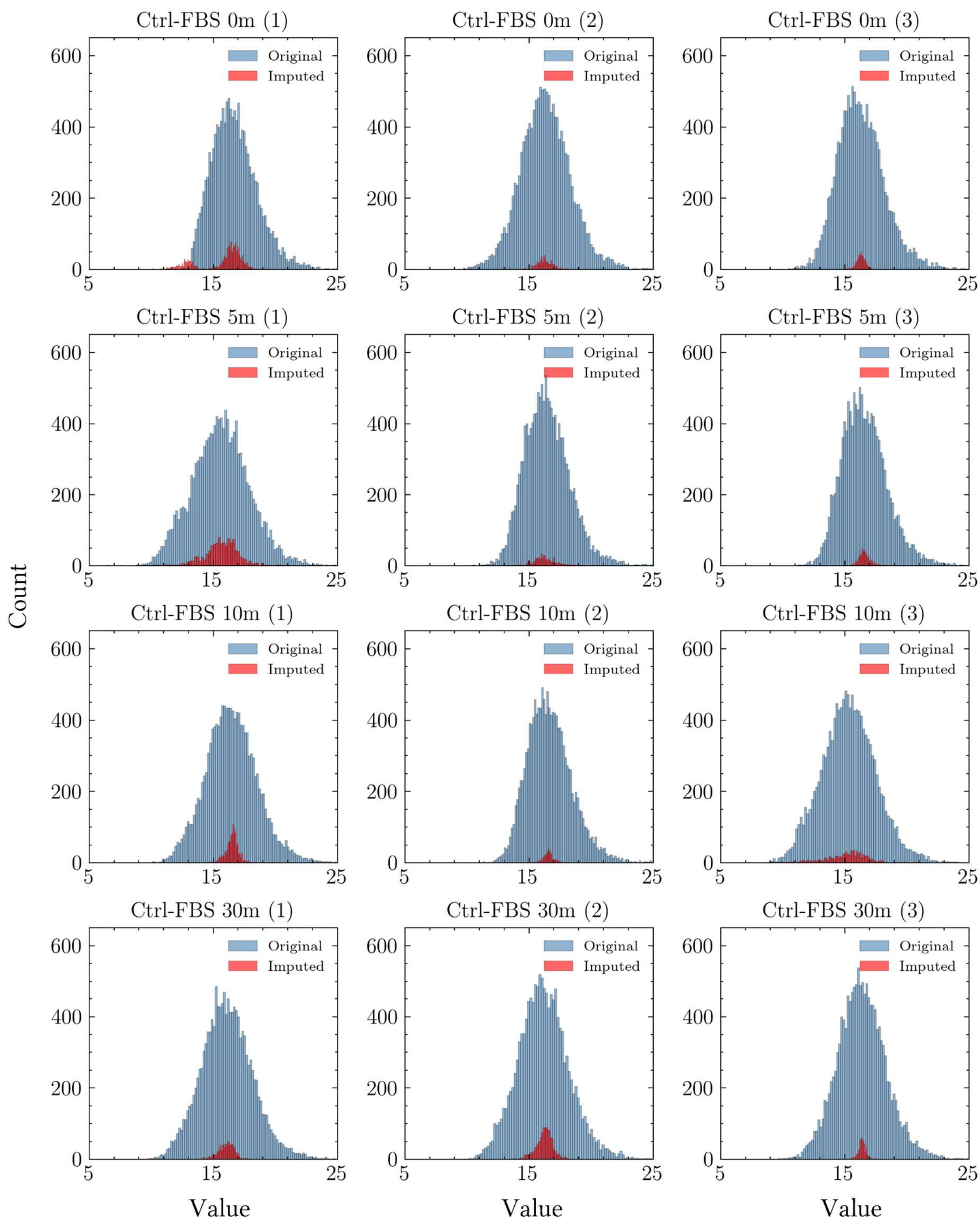


**Figure S20:** Histogram analysis of the frequency distribution of BPCA imputed phosphoproteome data Control minus FBS.

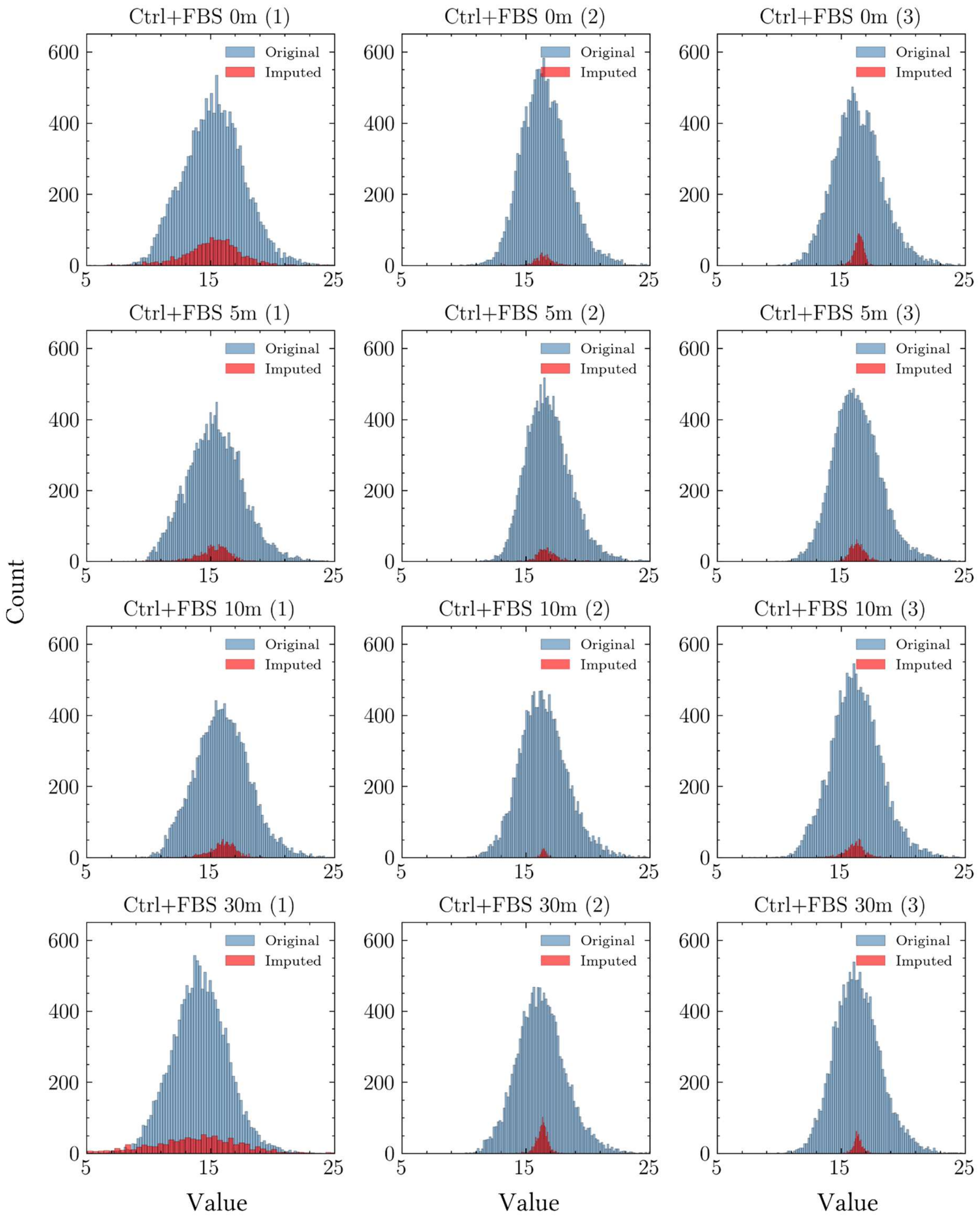


**Figure S21:** Histogram analysis of the frequency distribution of BPCA imputed phosphoproteome data Control plus FBS.

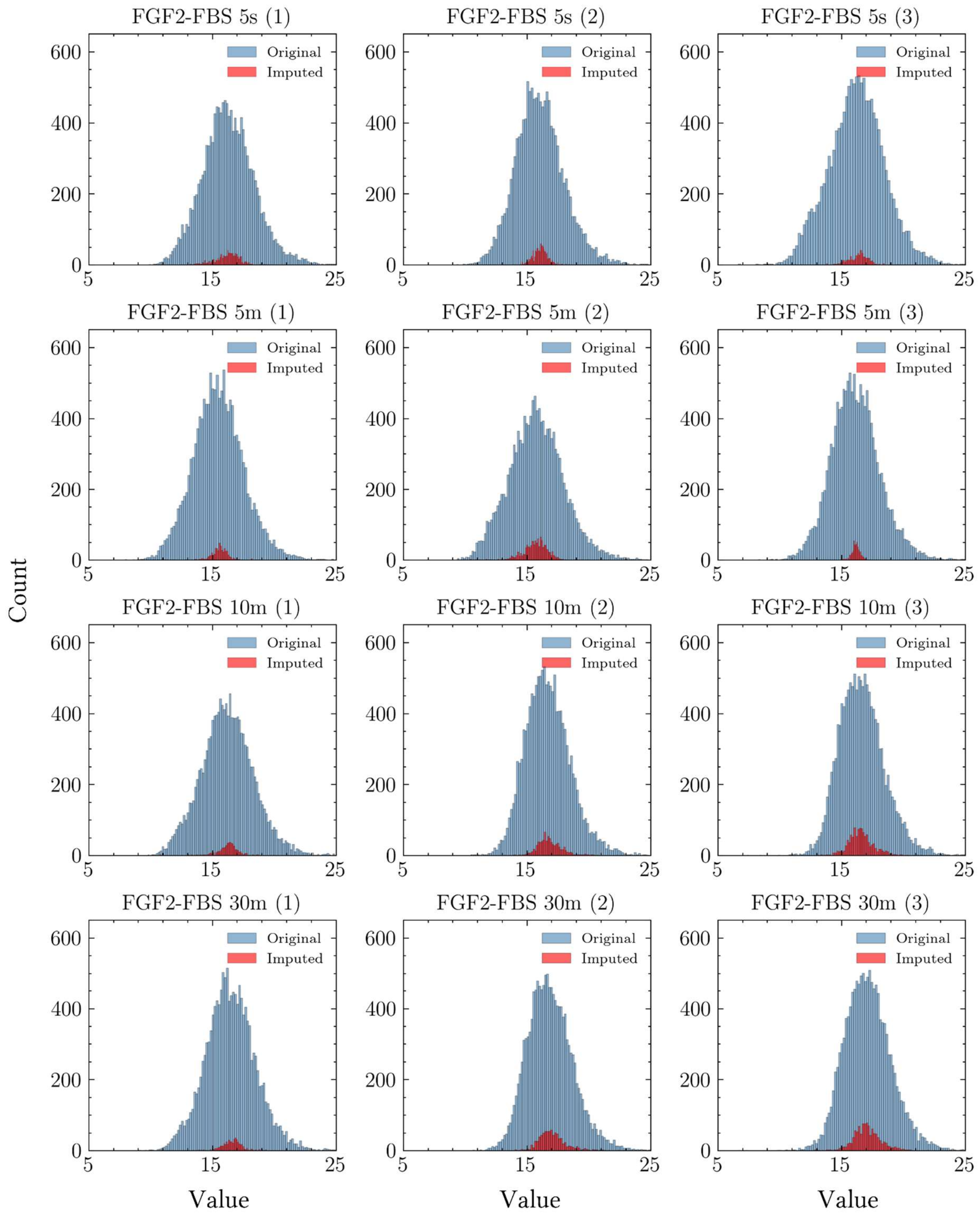


**Figure S22:** Histogram analysis of the frequency distribution of BPCA imputed phosphoproteome data FGF2 minus FBS.

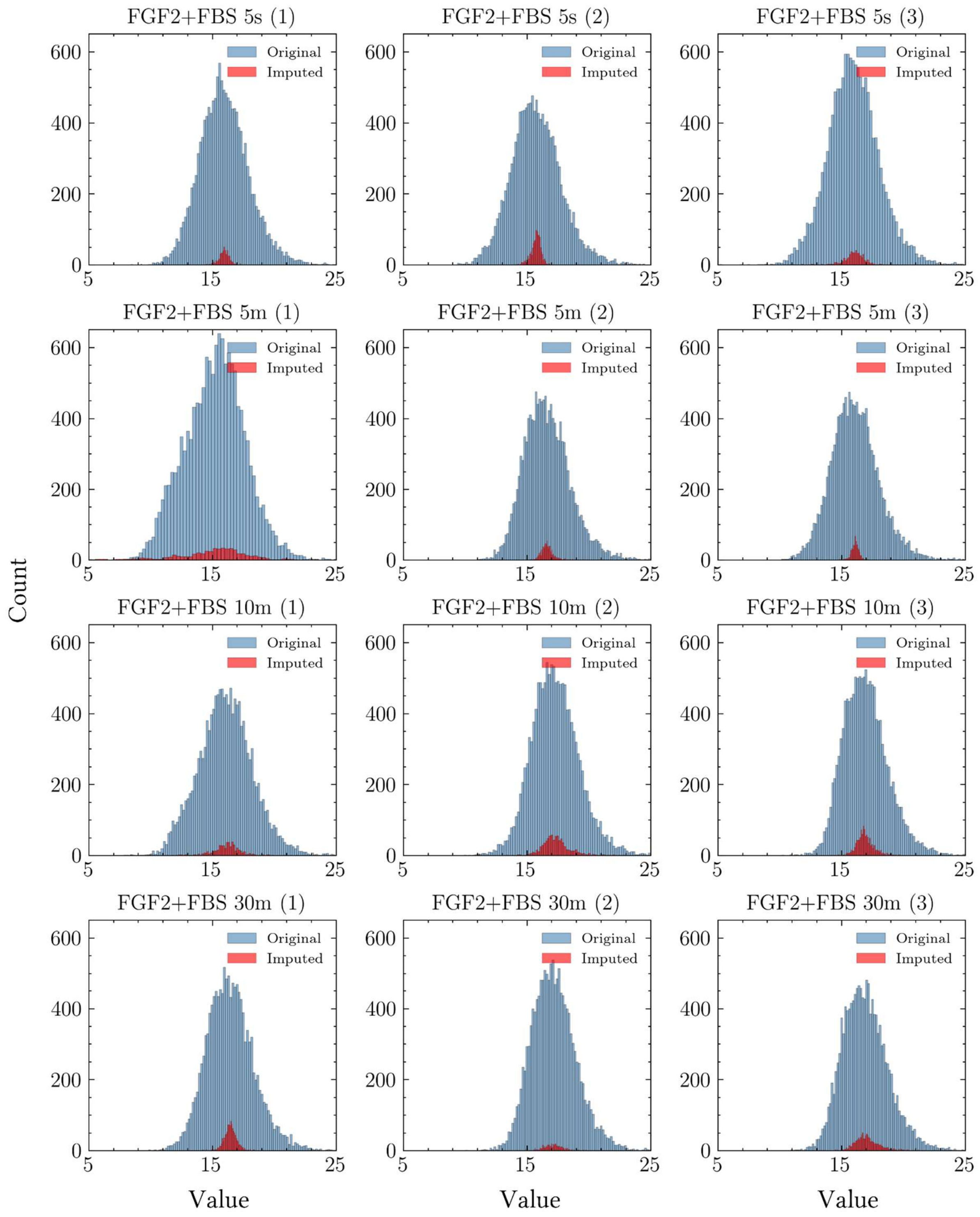


**Figure S23:** Histogram analysis of the frequency distribution of BPCA imputed phosphoproteome data FGF2 plus FBS.

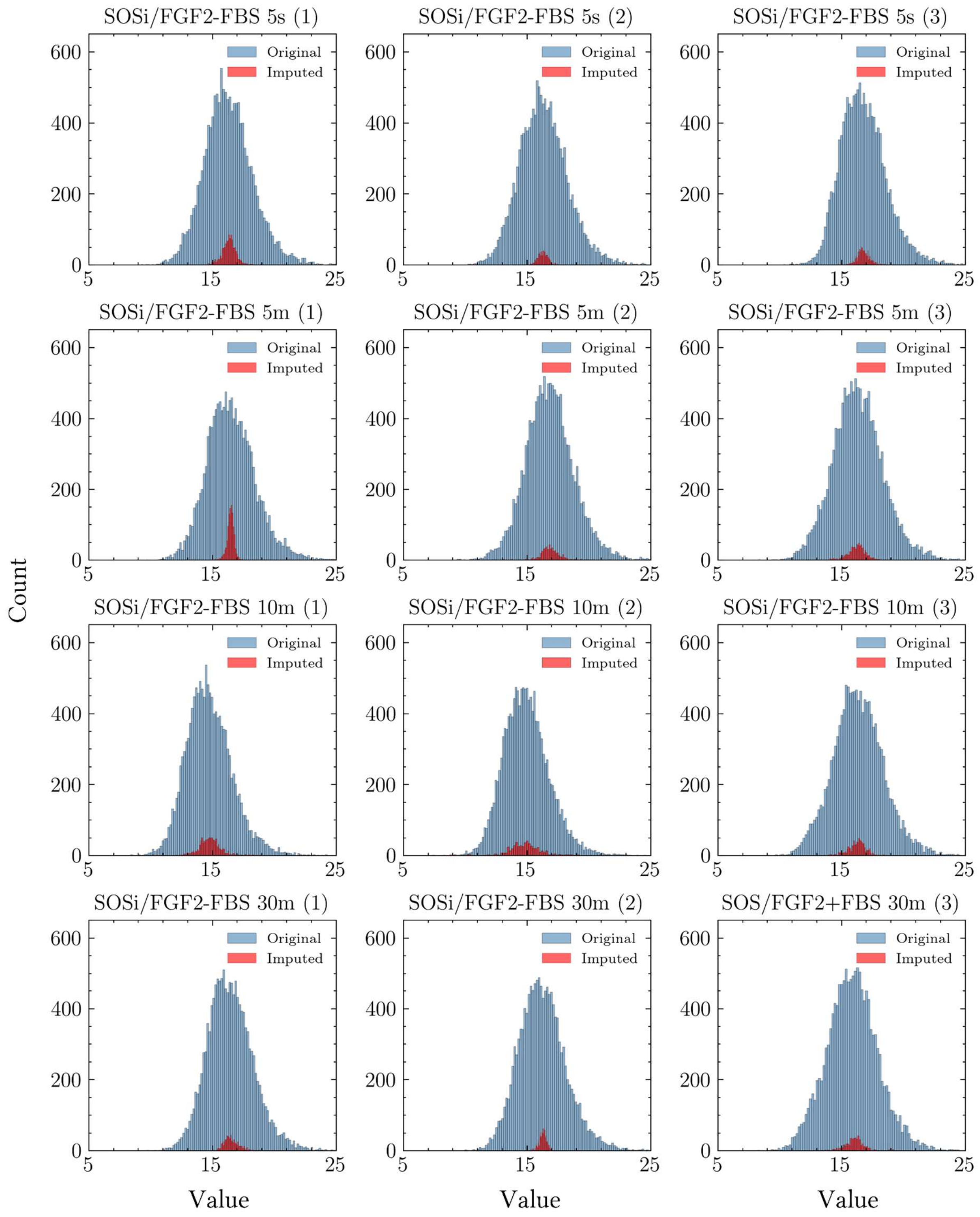


**Figure S24:** Histogram analysis of the frequency distribution of BPCA imputed phosphoproteome data SOSi/FGF2 minus FBS.

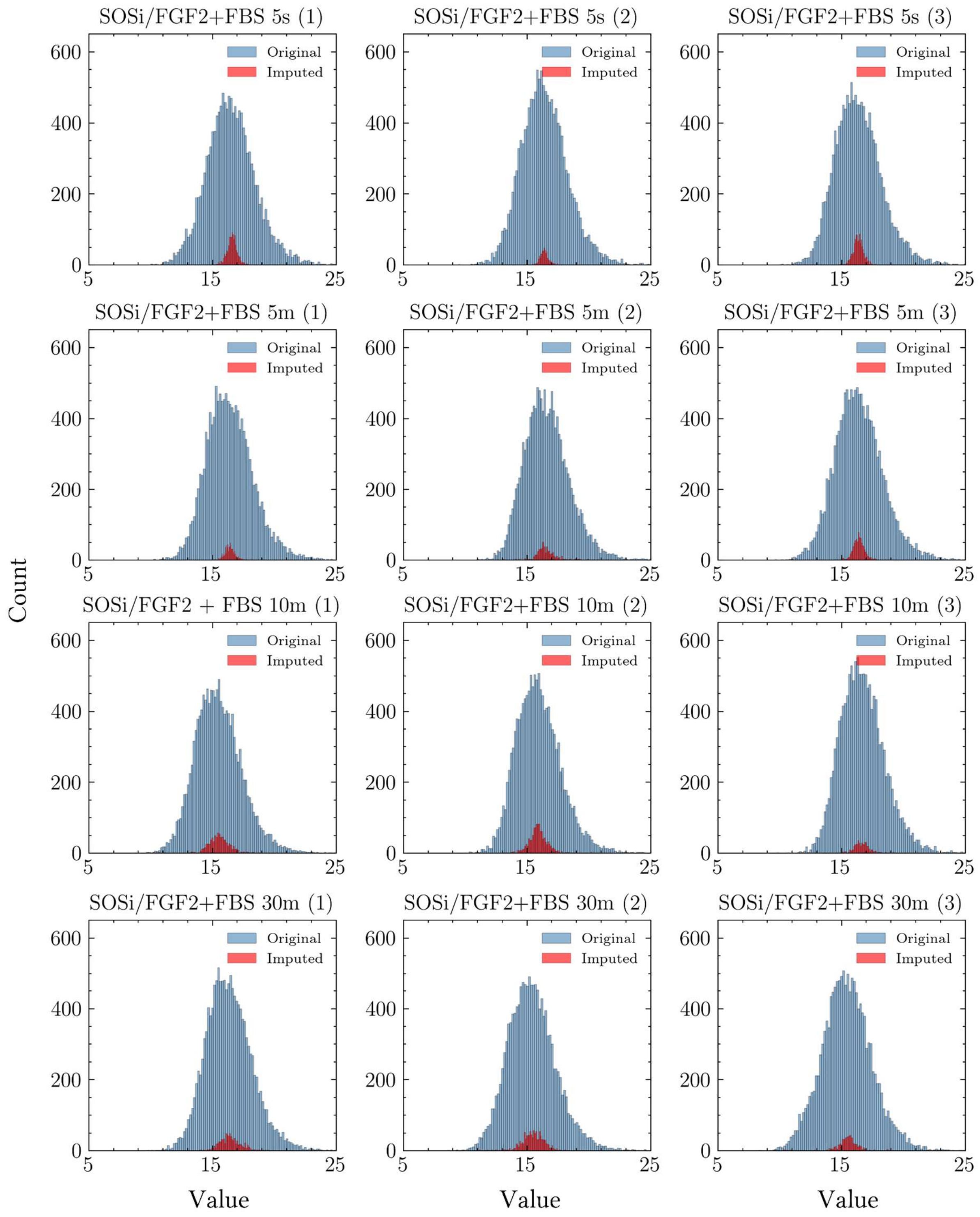


**Figure S25:** Histogram analysis of the frequency distribution of BPCA imputed phosphoproteome data SOSi/FGF2 plus FBS.

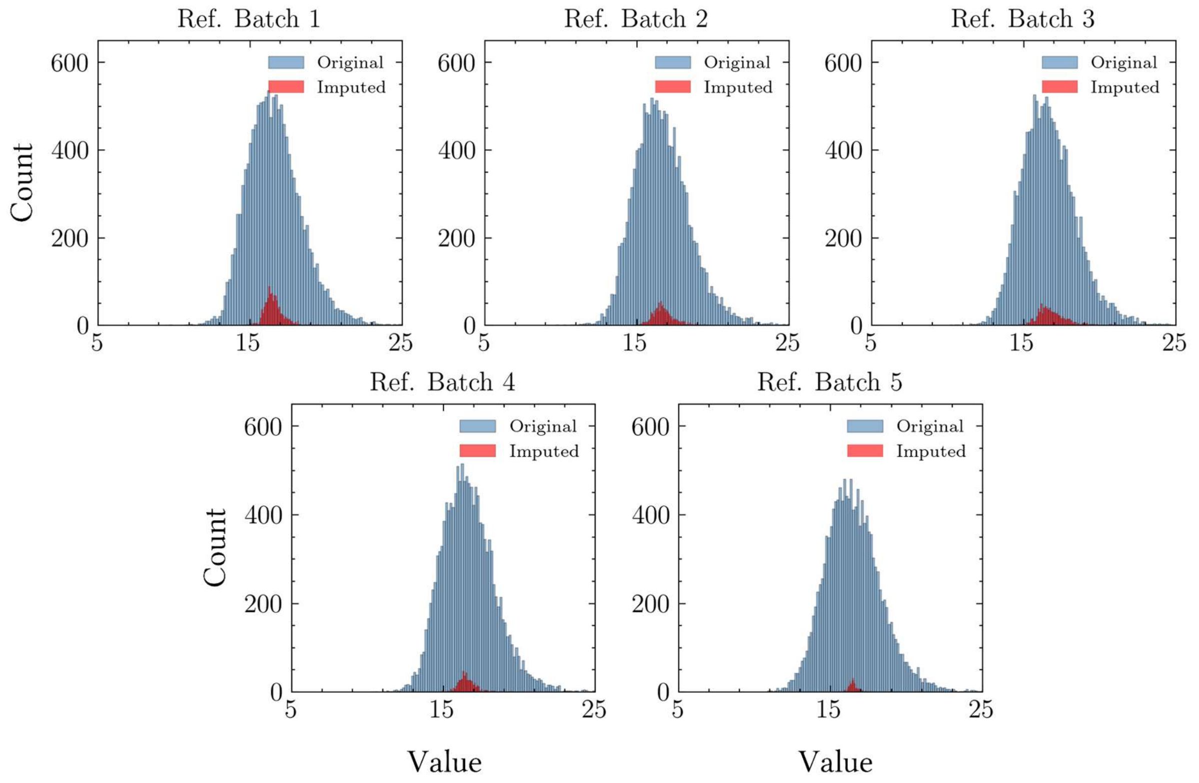


**Figure S26:** Histogram analysis of the frequency distribution of BPCA imputed phosphoproteome data Reference Samples.
